# A single-cell atlas linking intratumoral states to therapeutic vulnerabilities across cancers

**DOI:** 10.64898/2026.02.18.706316

**Authors:** María González-Bermejo, Laura Serrano-Ron, Santiago García-Martín, Óscar Lapuente-Santana, Ignacio Sanz Portillo, Pablo González-Martínez, Gonzalo Gómez-López, Fátima Al-Shahrour

## Abstract

Intratumoral heterogeneity (ITH) is a major determinant of therapeutic failure, yet its impact on drug response across cancers remains incompletely understood. Here, we present the Therapeutic Cancer Cell Atlas (TCCA), a pan-cancer single-cell resource integrating ∼1.8 million transcriptomes from 537 patients and 183 cancer cell lines spanning 34 tumor types. By combining single-cell transcriptomics with copy-number alteration inference and computational drug-response prediction, we systematically map therapeutic heterogeneity at subclonal resolution across cancers. Using this framework, we identify ten recurrent therapeutic clusters that capture conserved and context-specific drug vulnerabilities across tumor lineages. Notably, therapeutic heterogeneity is largely decoupled from genomic and transcriptomic diversity and instead arises from distinct functional transcriptional programs and tumor microenvironment (TME) states. Integration with transcriptional metaprograms and TME archetypes reveals how stress responses, proliferative states, lineage programs, and immune context shape drug sensitivity beyond tissue of origin. We further demonstrate the translational relevance of TCCA by linking therapeutic clusters to patient outcomes and validating predicted vulnerabilities using pharmacogenomic datasets, including clinically actionable examples in aggressive tumor subtypes. Together, TCCA provides a multidimensional atlas connecting subclonal states, microenvironmental context, and drug response, offering a scalable framework to guide therapeutic prioritization, drug repurposing, and combination strategies in precision oncology.

## Introduction

Tumor heterogeneity is a fundamental hallmark of cancer, with profound implications for disease progression, therapeutic resistance, and adverse clinical outcomes. This heterogeneity can be broadly categorized into intertumoral heterogeneity, referring to differences between tumors across or within patients, and intratumoral heterogeneity (ITH), which describes the coexistence of genetically and phenotypically distinct cell populations within a single tumor. ITH may manifest spatially, across different tumor regions, or temporally, as tumors evolve over time^1^. High levels of ITH and ongoing clonal evolution have been consistently associated with poor prognosis and reduced treatment efficacy, underscoring their clinical relevance^2^.

The origins of ITH lie in the dynamic interplay between genomic and epigenomic alterations, transcriptional plasticity, and interactions with the tumor microenvironment (TME), which together shape tumor development, progression, and response to therapy^2,3^. Advances in single-cell technologies, particularly single-cell RNA sequencing (scRNA-seq), have enabled high-resolution dissection of tumor ecosystems, revealing diverse malignant states and complex interactions between tumor and non-malignant compartments^4^. Pan-cancer single-cell atlases have shown that transcriptional programs underlying ITH often recur across tumors and cancer types, suggesting conserved functional states that transcend tissue of origin^5–9^.

Despite these advances, most existing single-cell atlases have focused on characterizing transcriptional and cellular diversity, with limited attention to the therapeutic consequences of ITH. As a result, the relationship between subclonal functional states and drug response remains poorly defined, constraining the translational potential of these resources. This gap is particularly relevant given growing evidence that tumors with similar genetic profiles can exhibit markedly different therapeutic sensitivities, reflecting functional plasticity and context-dependent dependencies rather than mutation burden alone^10^.

Recent computational approaches have begun to infer drug responses directly from single-cell transcriptomic data, enabling systematic prediction of therapeutic vulnerabilities at cellular resolution. However, these tools have largely been applied in isolated contexts, and a pan-cancer framework linking inferred drug responses to subclonal architecture, transcriptional programs, and the TME is still lacking. Moreover, the extent to which therapeutic heterogeneity aligns or diverges with genomic and transcriptional heterogeneity across cancers remains unresolved.

To address these challenges, we constructed the Therapeutic Cancer Cell Atlas (TCCA), a pan-cancer single-cell resource derived from publicly available scRNA-seq datasets comprising ∼1.8 million cells from 537 cancer patients and 183 cancer cell lines across 34 tumor types. By integrating single-cell transcriptomes with copy-number alteration inference^15^ and drug-response predictions^11,14^, we establish a unified framework to interrogate subclonal architecture, therapeutic heterogeneity, transcriptional programs, and TME archetypes within and across tumors. Using this approach, we identify recurrent pan-cancer therapeutic clusters that capture conserved and context-specific drug vulnerabilities at subclonal resolution. We show that therapeutic heterogeneity is largely decoupled from genomic and transcriptomic diversity and instead emerges from distinct functional states and microenvironmental contexts. Together, these findings position TCCA as a multidimensional atlas that connects intratumoral heterogeneity to actionable therapeutic insights, providing a foundation for functional precision oncology across cancer types.

## Results

### The TCCA is a harmonized single-cell resource for pan-cancer therapeutic analysis

The Therapeutic Cancer Cell Atlas (TCCA) was assembled from 36 public scRNA-seq studies (Supplementary table 1) to enable systematic investigation of therapeutic responses and recurrent drug sensitivity patterns across tumour types. The TCCA integrates curated clinical annotations with single-cell profiles spanning ∼1.8M cells across 34 tumor types, enabling stratified analyses across diverse clinical variables. All datasets were preprocessed through a unified pipeline to ensure consistent quality control, cell-type annotation, and cross-study comparability. This curated atlas serves as the basis for downstream analyses linking clonal architecture, gene expression, and therapeutic prediction (Fig. 1a).

**Figure 1.**
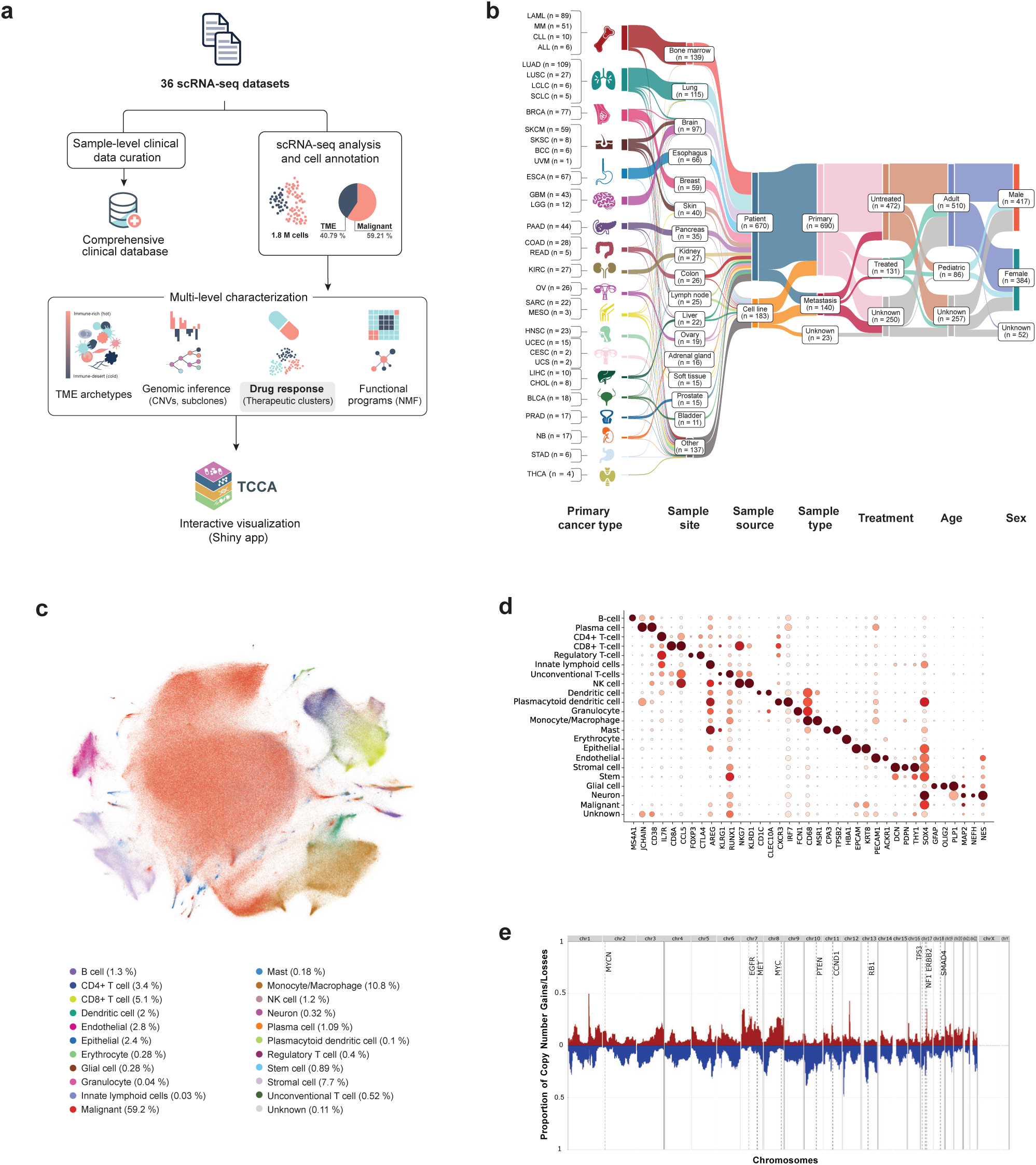
Pan-cancer overview of the Therapeutic Cancer Cell Atlas (TCCA). **a** Workflow integrating clinical and single-cell RNA-seq data, including TME archetypes, genomic inference, drug response prediction, and NMF-based functional analysis. **b** Sankey plot showing sample distribution by cancer type across key clinical variables. **c** UMAP embedding of the 1M malignant cells and 800k TME cells integrated across multiple studies, colored by cell type. **d** Validation of the cell-type annotation based on expression of canonical marker genes. **e** Pan-cancer CNV patterns in TCCA samples, with the y-axis showing the proportion of subclones carrying genomic alterations across chromosomes (x-axis).

For inclusion in the atlas, samples were required to contain at least 100 malignant cells and matched clinical metadata. After filtering, the final cohort comprised primary (n = 690) and metastatic (n = 140) tumors, treated (n = 131) and untreated (n = 472) samples, pediatric (n = 86) and adult (n = 510) patients, with balanced representation of both sexes (417 males, 384 females) (Supplementary table 1). Lung adenocarcinoma (LUAD), acute myeloid leukemia (LAML), breast cancer (BRCA), and esophageal carcinoma (ESCA) were the most represented tumor types (Fig. 1b).

Malignant and non-malignant compartments were identified using copy-number profiles inferred with SCEVAN^15^ (See Methods), yielding 1,089,024 malignant and 750,218 non-malignant cells. These cells were integrated across studies using GPU-accelerated scANVI^16^, enabling robust cross-sample alignment. Across cancers, epithelial and immune compartments predominated (Fig. 1c). Lineage-specific clusters resulting from harmonization were further validated using well-known cell type-specific biomarkers (Fig. 1d). The full TCCA data matrix, including cell-type annotations and curated clinical metadata, is publicly available and can be interactively explored at https://tcca.bioinfo.cnio.es/.

### The TCCA reveals diverse subclonal genomic architectures underlying tumor heterogeneity

Inference of copy number variations (CNVs) identified malignant subclones across all tumor types, exposing extensive inter- and intra-tumoral heterogeneity. As expected, malignant cells exhibited a higher number of CNAs (Supplementary Fig. 1), including recurrent amplifications on 1q21-q22 and 12q13, and deletions on 12p13 regions harboring known cancer driver regions across epithelial and hematologic cancers^17–19^ (Fig. 1e). Tumor-type-specific CNA patterns were observed such as the gain of chromosome 7 and loss of chromosome 10 in glioblastoma (GBM)^20^, 3p loss in kidney renal clear cell carcinoma (KIRC)^21^, and distinct CNV patterns between LUSC and LUAD on chromosomes 3, 5, and 8^22^ (Supplementary Fig. 2). Subclonal complexity varied markedly across cancer types (Supplementary Fig. 3). Subclonal burden did not differ by sex but increased with patient age. Primary tumors harbored more subclones per 1,000 cells than metastatic samples, consistent with metastatic bottlenecks^23^, and untreated tumors showed higher clonal diversity than treated ones, likely reflecting therapy-induced selection^24^ (Supplementary Fig. 4a-d).

### Drug response landscape at the subclonal level reveals recurrent pan-cancer therapeutic clusters and context-specific vulnerabilities

To map therapeutic heterogeneity at the subclonal level, we applied scTherapy^14^, using differential expression between malignant and non-malignant cells to predict drug efficacy for 3,695 compounds. On average, 158 drugs were predicted to exhibit high or moderate efficacy per subclone (Supplementary Fig. 5). Clustering subclones by shared drug-response profiles revealed ten recurrent pan-cancer therapeutic clusters (TCs) (Fig. 2a, Supplementary Fig. 6). Analysis of the distribution of cancer types across therapeutic clusters revealed that most TCs comprise subclones from multiple tumor lineages, underscoring the existence of recurrent, pan-cancer therapeutic programs (Supplementary Fig. 7). A subset of TCs, however, displayed strong tissue specificity, for example, TC5 and TC6 were dominated by gliomas (Fisher’s exact test, OR = 29.4, p < 2.2×10^−16^) and acute myeloid leukemia (OR = 25.4, p < 2.2×10^−16^), respectively indicating that certain vulnerabilities are tumor-dependent. This pattern highlights the coexistence of conserved and lineage-restricted therapeutic states across cancers. CNV profiling further revealed cluster-specific patterns of chromosomal instability, while previously identified recurrent events were shared across multiple TCs(Fig. 2d, Supplementary Fig. 8), reinforcing their role as conserved genomic drivers of drug sensitivity patterns. Across clinical contexts, metastatic samples were enriched in TC3 and TC5, whereas untreated tumors were biased toward TC1 and TC3. Pediatric and young adult tumors were overrepresented in TC6 and TC8-9, while adult and elderly samples showed broader distributions (Supplementary Fig. 9a-c), indicating that therapeutic heterogeneity reflects both tumor evolution and treatment history.

**Figure 2.**
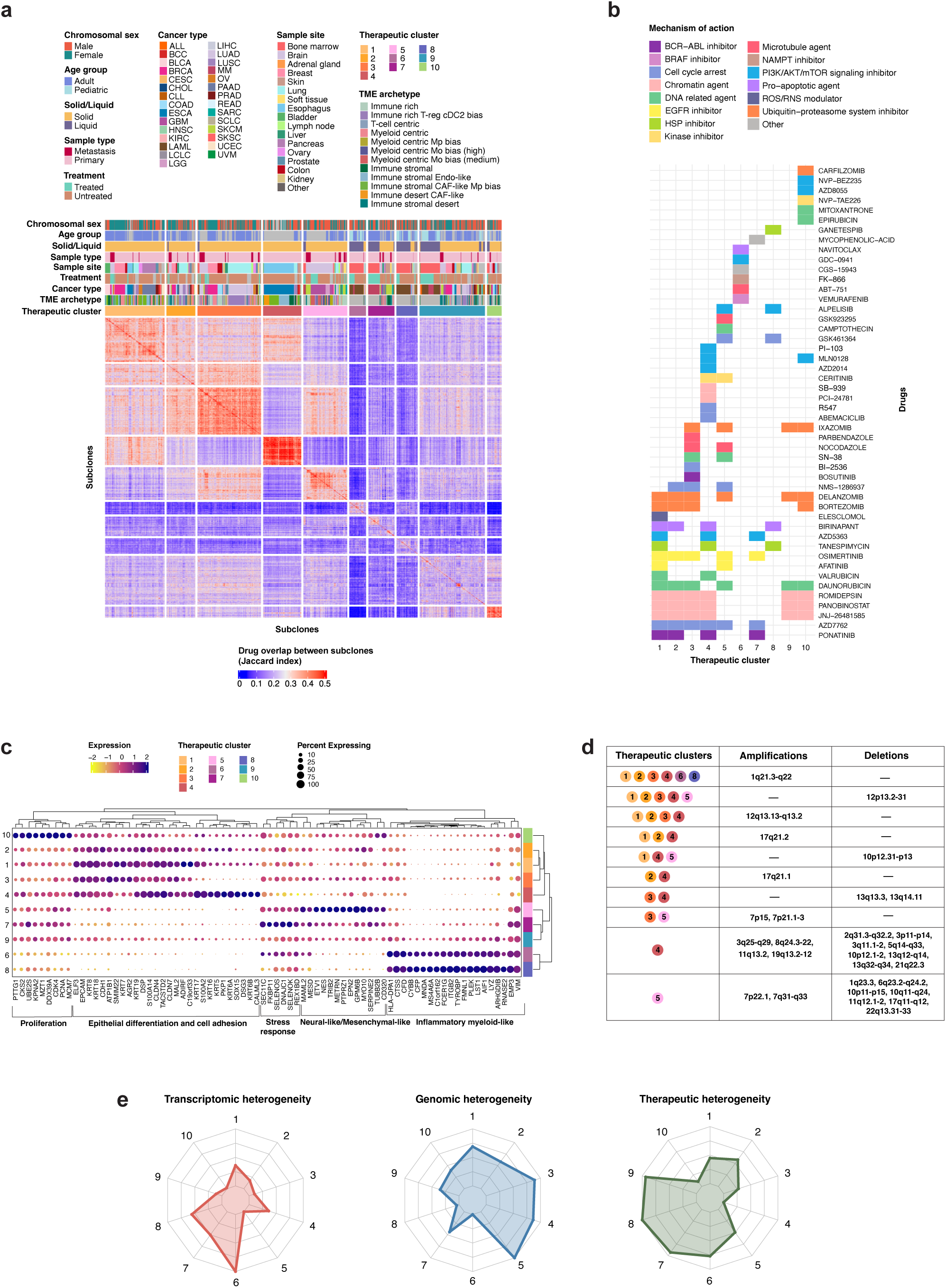
Pancancer therapeutic clusters and their association with molecular and microenvironmental features. **a** Heatmap of pairwise drug prediction overlaps (Jaccard index, color gradient) between genomic subclones. Only high-confidence malignant cells were used (611 samples, 2,627 subclones, 637,062 cells; see Methods). Spectral clustering identified 10 therapeutic clusters. Top annotations indicate sample characteristics, including sex, age, sample site, primary/metastasis status, solid/liquid tumor type, treatment condition, cancer type, TME archetype, and therapeutic cluster. **b** Tile plot of drugs predicted by scTherapy (y-axis) in ≥60% of subclones per therapeutic cluster (x-axis). Tiles are colored by mechanism of action. **c** Dot plot of cluster marker genes identified via differential expression (FindAllMarkers, Wilcoxon rank-sum test; positive markers only, avg_log2FC ≥ log2(1.5), pct.1 ≥ 0.5, pct.1 - pct.2 ≥ 0.2). Color indicates expression level; dot size represents the proportion of cells expressing the gene. **d** Table of CNVs (amplifications and deletions) present in ≥50% of subclones within each therapeutic cluster. Some CNVs are shared across multiple clusters, which are indicated in the therapeutic cluster column. **e** Radar chart showing heterogeneity of each therapeutic cluster across three dimensions: transcriptional (mean Shannon entropy of log-normalized gene expression per cell), therapeutic (mean pairwise Jaccard distance between subclone drug predictions), and genomic (mean Shannon entropy of subclone copy number profiles across cytobands). Values are scaled relative to the maximum observed across clusters, allowing direct comparison of multidimensional.

Three closely related clusters (TC1-3) shared vulnerabilities and were enriched in pancreatic, breast, lung, bladder, esophageal, uterine, and prostate cancers (Fig. 2a). These TCs exhibited sensitivity to chromatin-modifying agents, including HDAC inhibitors such as romidepsin and panobinostat. This responsiveness to HDAC inhibition is consistent with recurrent alterations in *KRAS*, *ERBB2*, and *BRAF*^25,26^, key MAPK pathway drivers commonly found in the cancer types enriched within these clusters, which can modulate sensitivity to HDAC inhibitors^27^. Collectively, TC1-3 displays overlapping sensitivity to epigenetic, proteasome, apoptotic, and signaling-targeted agents, defining a shared vulnerability axis across epithelial tumors (Fig. 2b). Transcriptomic markers highlighted epithelial identity and adhesion (*EPCAM*, *KRT8*, *CLDN4*) alongside oncogenic drivers (*S100A14*, *TACSTD2*) (Fig. 2c). Genomically, they are characterized by amplifications at 12q13.13-q13.2 encompassing *KRT* genes and oncogenes (*MDM2*, *CDK4*, *STAT6*, *DDIT3*, *GLI1*), a recurrent aberration observed in multiple epithelial cancers including NSCLC^28^ and sarcomas^29^ (Fig. 2d). TC4, enriched in esophageal cancer subclones (Fig. 2a), is a therapeutically homogeneous cluster exhibiting shared sensitivities to chromatin modifiers, cell cycle and kinase inhibitors, and PI3K/AKT/mTOR and EGFR pathway inhibitors, and is marked by recurrent 3q amplifications and 5q deletions typical of esophageal squamous cell carcinoma^30^ (Fig. 2d). TC5, predominantly composed of gliomas and melanoma-derived brain metastases, revealed shared therapeutic and molecular features between primary brain tumours and brain metastasis (Fig. 2a). Despite their distinct origins, TC5 subclones converged on vulnerabilities targeting cell cycle regulation, DNA replication, and PI3K signaling, consistent with frequent PI3K hyperactivation in brain tumours^31^ (Fig. 2b). TC5 showed high genomic and transcriptional heterogeneity and displayed recurrent deletions affecting immune-regulatory and signaling pathways, consistent with aggressive brain tumor biology(Fig. 2e)^33–39^. Shared deletions with other TCs, including 12p13.2-p13.31 encompassing *BCL2L14,* KLRC2/3/4, CLEC family genes, and *ETV6,* highlighted common vulnerabilities in apoptotic regulation and transcriptional repression^40,41^ (Fig. 2d). Transcriptomic profiling showed co-activation of neural and mesenchymal programs (*NES*, *TUBB2B*, *MYO10*), indicative of a hybrid neuro-mesenchymal phenotype supporting invasiveness and plasticity (Fig. 2c).

TC6-TC9 showed the highest therapeutic heterogeneity, reflecting divergent subclonal dependencies on proteostasis and stress-response pathways (Fig. 2e, Supplementary Fig. 10a-b). TC6 and TC8, enriched in LAML, were sensitive to apoptotic, metabolic, and HSP90-targeting agents, whereas TC7 and TC9 converged on proteasome and ER-stress vulnerabilities across solid and hematologic malignancies^42–45^. These clusters lacked recurrent CNAs, indicating that functional stress states rather than genomic alterations underlie their drug sensitivities.

Overall, comparative analysis of TCs revealed pronounced differences in both intra- and inter-cluster diversity. Although genomic and transcriptomic heterogeneities were moderately negatively correlated (r = −0.48), therapeutic heterogeneity showed little association with either molecular feature alone, suggesting a multilayer molecular basis for drug-response diversity. Some clusters, such as TC5, displayed high transcriptional and genomic variability but only modest therapeutic divergence, whereas others (TC6-TC9) exhibited broad therapeutic heterogeneity despite relatively stable molecular features (Fig. 2e). Importantly, TCs with high therapeutic heterogeneity are more likely to harbor subclones with divergent drug sensitivities, complicating treatment strategies and requiring combination or adaptive approaches. In contrast, TCs with low therapeutic heterogeneity, such as TC4 and TC10, may be more uniform for therapeutic targeting (Supplementary Fig. 10). Together, this integrated map highlights the complex but structured nature of subclonal therapeutic heterogeneity, revealing conserved dependencies that transcend tissue of origin and may inform combination strategies in precision oncology.

### Systematic investigation of therapeutic heterogeneity across multiple cancer types

Analysis across tumor types revealed marked variability in the degree and structure of therapeutic heterogeneity. Cancers such as breast cancer (BRCA) and lung adenocarcinoma (LUAD) exhibited broad therapeutic diversity, with subclones spanning multiple TCs, whereas other tumor types showed more homogeneous profiles restricted to one or two dominant clusters (Fig. 3a). Despite this diversity, several cancers were strongly enriched for specific TCs, including esophageal cancer (ESCA; TC4), glioblastoma and low-grade glioma (GBM/LGG; TC5), and bladder cancer (BLCA; TC1), indicating that for certain tumors, specific treatments are likely to be broadly effective across most subclones (Fig. 3b). Cervical carcinoma (CESC), uveal melanoma (UVM), and chronic lymphocytic leukemia (CLL) displayed the highest therapeutic heterogeneity, whereas ESCA, head and neck squamous carcinoma (HNSC), and colorectal cancer (COAD) exhibited more uniform drug-response profiles (Fig. 3c).

**Figure 3.**
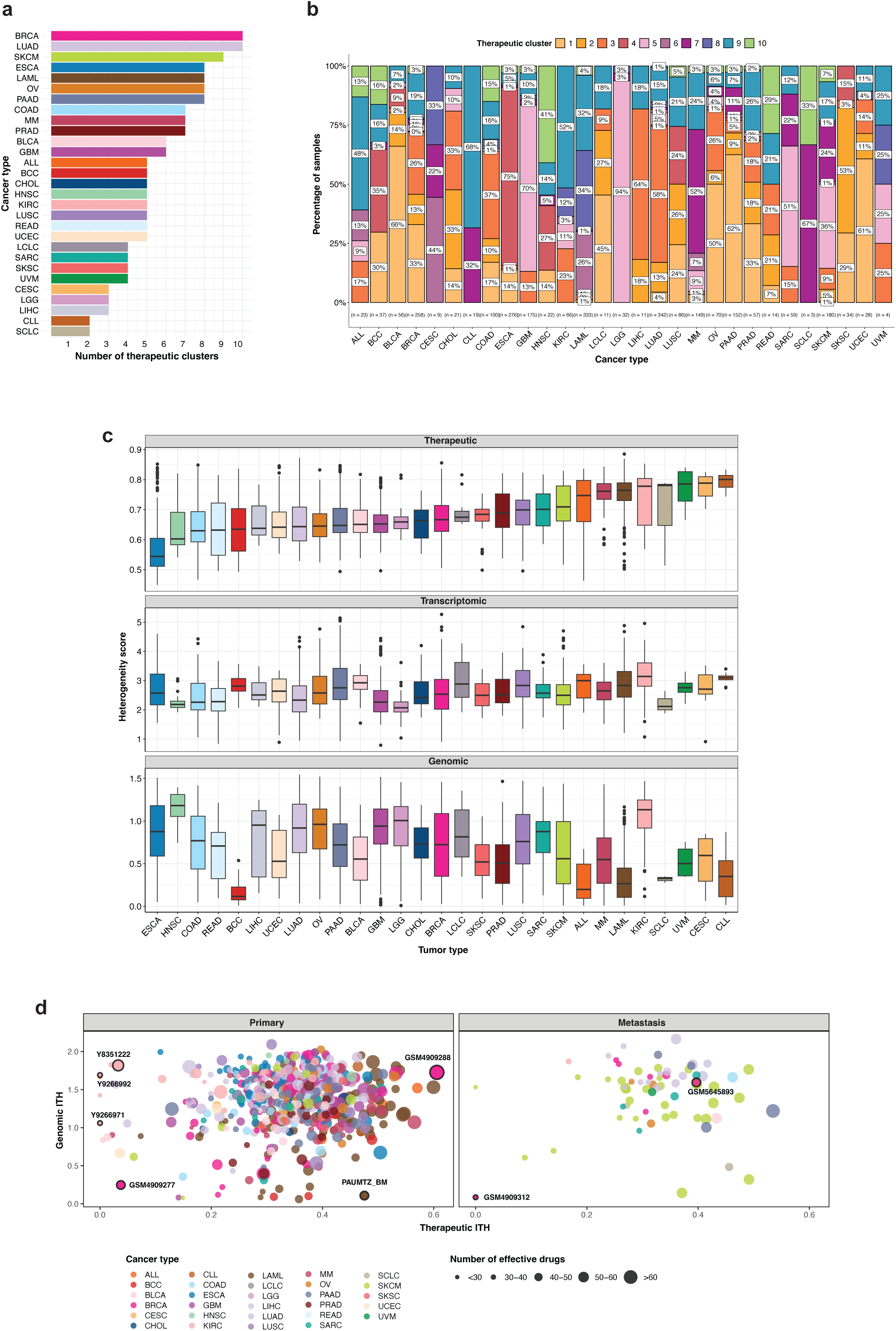
Cancer types are differentially distributed across therapeutic clusters. **a** Barplot illustrating, for each cancer type, the number of therapeutic clusters in which its subclones are represented. **b** Distribution of the 10 therapeutic clusters (TCs) across cancer types. Stacked bars represent the percentage of subclones assigned to each TC within each cancer type (n total = 2627 subclones). Colors indicate different TCs and numbers below each bar denote the total number of subclones in each cancer type. **c** Quantitative analysis of therapeutic, transcriptomic, and genomic heterogeneity of subclones across cancer types. Boxplots show heterogeneity scores for each subclone across cancer types (each point represents a subclone). Cancer types on the x-axis are ordered according to increasing therapeutic heterogeneity. **d** Correlation between genomic and therapeutic ITH sample-wise. In breast cancer, patients with low genomic and therapeutic heterogeneity (e.g., GSM4909277) were dominated by TC1-TC2 subclones and exhibited limited predicted drug sensitivity (42 drugs). In contrast, highly heterogeneous tumors (e.g., GSM4909288) contained additional TC8-TC9 subclones and showed broader drug responsiveness (77 drugs), including standard-of-care agents such as abemaciclib, alpelisib, epirubicin, lapatinib, paclitaxel, and docetaxel, as well as 28 drugs approved for other malignancies, underscoring repurposing potential. In hematologic cancers, a LAML patient (e.g., PAUMTZ_BM) with high therapeutic but low genomic heterogeneity showed sensitivity to 47 drugs, including daunorubicin (AML standard-of-care) and ponatinib (BCR-ABL inhibitor approved for CML/ALL), consistent with the enrichment of TC8–TC9 subclones. In contrast, KIRC patients (e.g., Y9266971, Y9266992, Y8351222) exhibited variable genomic heterogeneity with minimal therapeutic diversity. Cases with higher genomic heterogeneity showed an expanded set of effective drugs (50 vs. 21 in low therapeutic heterogeneity samples), including sunitinib approved for KIRC and exclusively associated with TC9 indicating that increasing genomic complexity may expand druggable vulnerabilities.

The TCCA further enables systematic evaluation of predicted therapeutic responses in relation to drug approval status. Of the 158 compounds predicted to be effective across subclones, 54 (34%) are clinically approved, including 39 (72%) approved for oncology indications, while 58 (37%) are currently under clinical investigation (Methods). Notably, many FDA-approved agents showed predicted efficacy outside their current indications, highlighting substantial opportunities for therapeutic repurposing enabled by functional, state-based stratification. For example, subclones from KIRC were predominantly enriched in TC9 and exhibited consistent predicted sensitivity to proteasome inhibitors, including bortezomib and ixazomib (Supplementary Fig. 7). Although these agents are clinically approved for hematologic malignancies, their predicted efficacy in KIRC subclones highlights a proteostasis dependency that is not apparent from tumor type alone^45^. This illustrates how TCCA enables the identification of actionable, non-canonical drug vulnerabilities and supports rational repurposing strategies based on subclonal functional states rather than histological classification. Patient-level analysis revealed that tumors with broader TC composition exhibited expanded predicted drug sensitivity, including both standard-of-care and repurposing opportunities (Fig. 3d). Primary tumors displayed a wide range of heterogeneity profiles, whereas metastatic lesions were generally less diverse, consistent with clonal bottlenecks during dissemination.

Together, these analyses reveal that therapeutic heterogeneity operates partly independently of genomic architecture, varying both across and within tumor types and uncovers how specific tumor states rather than subclonal complexity alone shape drug responsiveness. By enabling patient- and subclone-resolved mapping of drug sensitivities, the TCCA provides a framework for identifying dominant vulnerabilities, uncovering repurposing opportunities, and prioritizing therapies tailored to functional tumor states rather than bulk tumor classification.

### Transcriptional programs reveal functional patterns underlying therapeutic clusters

To uncover the transcriptional determinants of drug sensitivity, we applied non-negative matrix factorization (NMF) to malignant cells, identifying 43 recurrent metaprograms (MPs) grouped into 11 functional families encompassing cell cycle/DNA repair, epithelial-to-mesenchymal transition (EMT), inflammation, stress/hypoxia, and lineage-specific cells (Fig. 4a, Supplementary table 2). Many MPs recapitulated transcriptional modules reported in previous pan-cancer atlases, confirming conserved axes of malignant plasticity^8,9^, while a novel family of cellular plasticity programs (MP22-MP24) linked to stemness, epithelial remodeling, and active signaling involving CDC42, RICTOR, MYH9, and ACTN4 emerged as a unique feature of this dataset (Fig. 4b). Additionally, functional enrichment using MSigDB collections and already published gene signatures from cancer cell states^5–8^ revealed that tumors segregate into groups defined by proliferative, metabolic, mesenchymal, immune/stress, and DNA-damage response states (Supplementary Fig. 11). Metastatic and treated tumors were dominated by chromatin remodeling, EMT, stemness, hypoxia, and stress hallmarks, whereas primary and untreated tumors were enriched in proliferative and cell cycle programs. Solid tumors preferentially activated hypoxia and EMT, in contrast to liquid tumors that relied on proliferation, while pediatric tumors clustered with developmental/stemness pathways distinct from adults.

**Figure 4.**
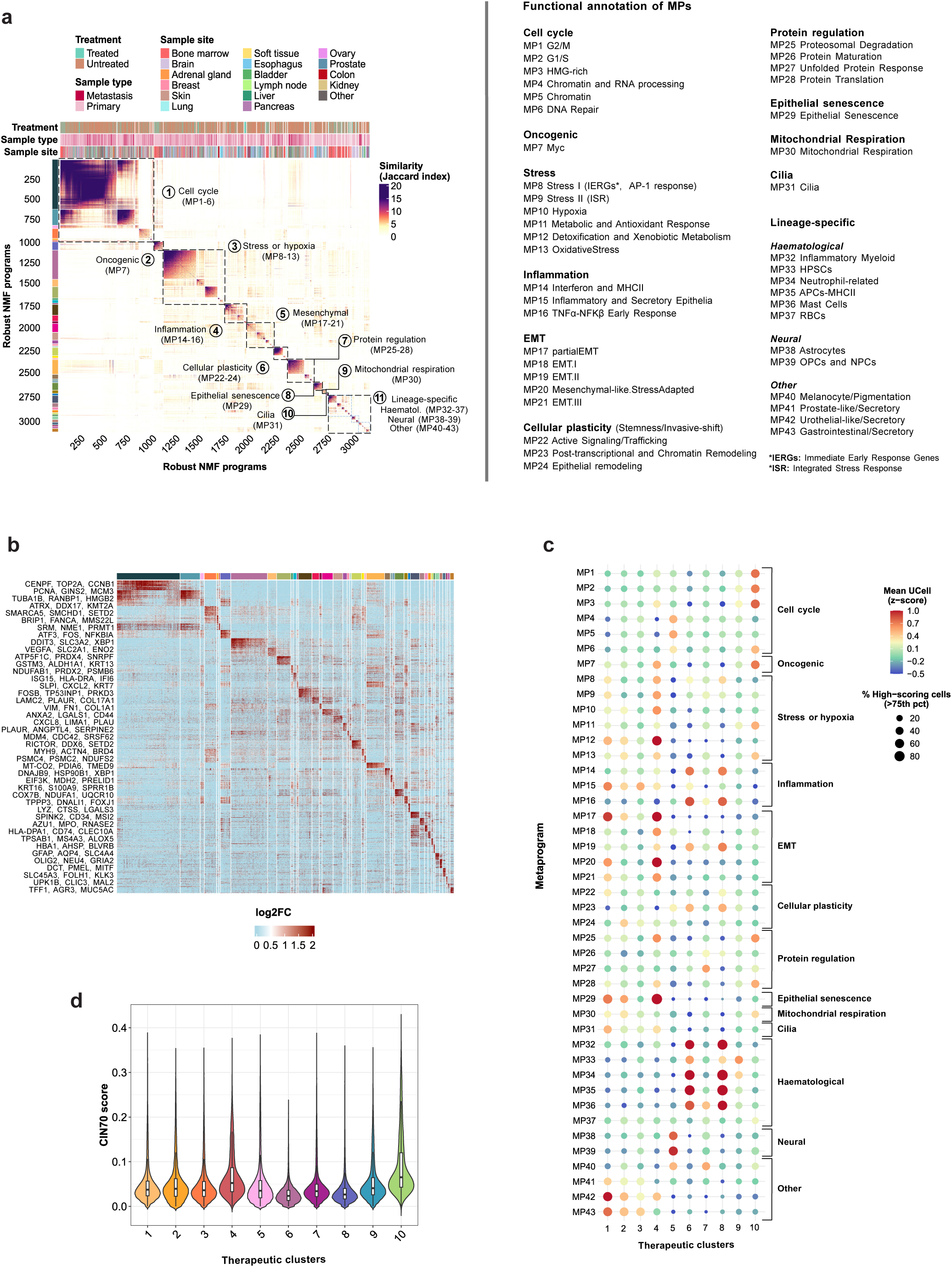
Mapping functional metaprograms to therapeutic clusters in TCCA. **a** Left: Heatmap of Jaccard similarity indices comparing 3,105 robust NMF programs based on their top 50 genes. Programs are clustered and grouped into 43 metaprograms (MPs; colored left annotation) and 11 MP families with related functions (black dashed lines, numbered). Right: List of all MP annotations grouped by family. **b** NMF scores for all MP genes (rows) across all robust NMF programs (columns), with representative genes are labeled. **c** Mean enrichment of MPs grouped by family (y-axis) across the 10 therapeutic clusters (x-axis). Color indicates z-scaled mean UCell score for cells in each TC, and dot size reflects the percentage of cells with high UCell scores (≥75th percentile within each MP). **d** Violin plot of CIN70 signature enrichment scores, estimating chromosomal instability across each therapeutic cluster.

Mapping MPs to TCs revealed distinct transcriptional determinants of drug responses (Fig. 4c). TC1 was enriched in EMT, inflammatory, and epithelial senescence programs, alongside lineage-specific urothelial and prostate signatures. This profile aligns with sensitivity to HDAC inhibitors, which reprogram cancer-associated fibroblasts by suppressing pro-inflammatory and pro-desmoplastic programs^46^, induce cellular senescence, and modulate EMT plasticity in a context-dependent mannerr^47^. TC2 and TC3 showed similar but more moderate activation of these modules. TC4 was enriched for MYC-driven and hypoxia/stress programs, aligning with mTOR pathway dependencies. TC5, composed mainly of gliomas and brain metastases, exhibited strong cell cycle and neural lineage signatures, supporting sensitivity to cell cycle inhibitors and microtubule-targeting agents. Hematologic-enriched TCs (6 and 8) showed inflammatory and myeloid lineage programs, while TC10 displayed proliferative MYC activation and protein regulation modules, matching its sensitivity to DNA-damaging and proteasome-targeting drugs. Notably, assessment of the CIN70 signature, a well-established marker of genomic instability^48^, further revealed significantly higher genomic instability in TC10 (Fig. 4d). To validate these associations, we correlated Beyondcell-derived^11^ drug sensitivity scores with MP enrichment (Supplementary Fig. 12). This analysis corroborated the TC-based findings, showing that cell cycle/DNA repair programs (MP1-MP6, enriched in TC5) correlated with sensitivity to DNA-damaging agents, PARP inhibitors, and microtubule-targeting drugs. The MYC program (MP7, dominant in TC4 and TC10) was associated not only with cell cycle and DNA-related agents but also with proteasome inhibitors and transcriptional regulators, consistent with MYC-driven proteotoxic stress arising from heightened biosynthetic demands. Stress and hypoxia programs (MP8-MP13, enriched in TC4) were linked to mTOR/PI3K inhibitor sensitivity, while inflammatory programs (MP14-MP16, active in TC1, TC6, and TC8) correlated with sensitivity to MAPK pathway modulators and enhanced immunotherapy response. Lineage-related MPs (MP32-MP43) revealed additional context-specific dependencies, such as sensitivity to EGFR and PI3K/mTOR pathway inhibitors in neural-like states (TC5) and MAPK kinase inhibitor responses in hematopoietic programs (TC6 and TC8).

### TME characterization in the TCCA cohort reveals new archetypes linked to drug response

We profiled 750,218 TME cells across 493 solid tumors from the TCCA cohort, recapitulating the five main TME archetypes described by Combes *et al.*^49^. This also revealed 7 novel immune and stromal subtypes, which we termed *immune-rich* Treg-cDC2 bias, *immune stromal CAF-like macrophage bias*, *immune stromal endo-like*, *myeloid-centric macrophage bias*, *myeloid-centric monocyte bias* (medium and high) and *immune-desert CAF-like* (Fig. 5a, Supplementary Fig. 13).

**Figure 5.**
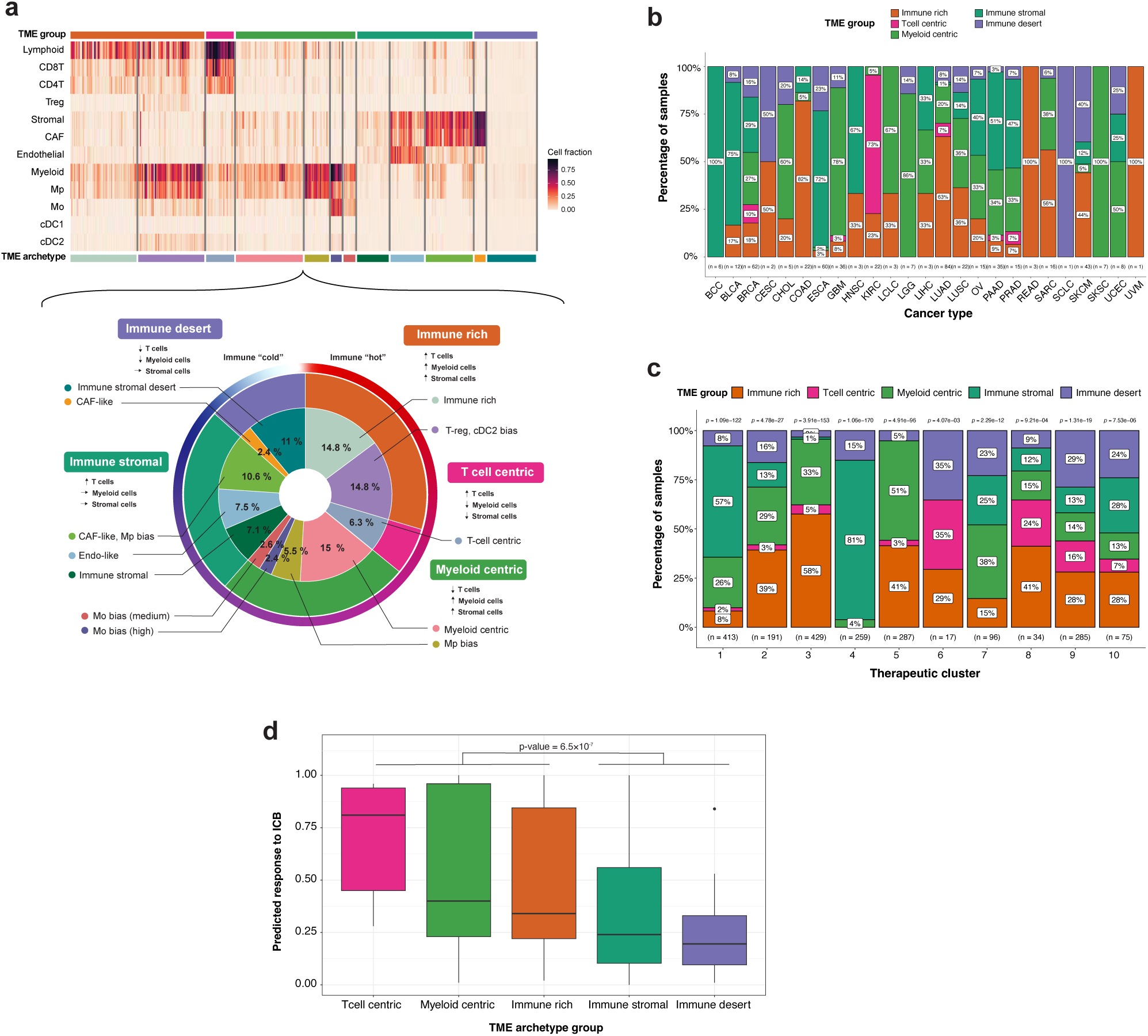
Tumor microenvironment archetypes capture conserved immune states across therapeutic clusters in TCCA. **a** Heatmap of 493 TCCA samples (excluding liquid tumors and samples with insufficient TME cells) clustered by the proportion of 10 immune and stromal cell types. The resulting clusters define distinct TME archetypes. The circular plot below orders TMEs from immune-hot (red) to immune-cold (blue); the outer ring depicts the five major TME groups, which correspond to those described by Combes and colleagues^49^, while the inner rings show their subdivisions into TME archetypes according to cell type biases. Some of this compositional biases differ from those reported by Combes^49^. **b** Distribution of TME groups across cancer types, where the y-axis represents the proportion of samples in each TME group. **c** Distribution of TME groups across therapeutic clusters; y-axis shows the proportion of subclones in each TME group. Numbers below each bar indicate the total number of subclones per TC, and numbers above show the Chi-squared p-value testing independence of TME distributions from the TCs. **d** Validation of TME archetypes using the Wang melanoma immune hot–versus–cold signature^51^. Switch Point (SP) scores, a measure derived from Beyondcell framework^11^, were computed per sample and summarized by TME group. The signature, composed of genes upregulated and downregulated in immune-hot relative to immune-cold tumors and previously validated in melanoma and glioblastoma, showed higher predicted immunotherapy sensitivity (1 − SP) in T cell–centric, myeloid-rich, and immune-rich samples, consistent with an immune-hot transcriptional profile.

TME composition varied substantially across tumor types (Fisher’s exact test, p = 1×10^−4^), with *immune-rich*, *immune-stromal*, and *myeloid-centric* archetypes widely distributed, while *immune-desert* and *T-cell-centric* archetypes were enriched in specific subtypes (Fig. 5b, Supplementary Fig. 14). Compared to primary tumors, metastatic lesions displayed enrichment of *immune-desert* TMEs (OR = 3.8, p < 1.9×10^−5^), depletion of *immune-stromal* (OR = 0.15, p < 1.1×10^−5^) and *T-cell-centric* archetypes (OR = 0.19, p = 0.07), and a modest increase in *immune-rich* TMEs (OR = 1.8, p = 0.034). These shifts likely reflect niche-specific immune remodeling during metastatic progression. Treatment status also influenced TME composition: untreated tumors showed a trend toward enrichment of *T-cell-centric* TMEs (OR = 4.01, p = 0.23) and were significantly enriched for *immune-stromal* states (OR = 2.37, p = 0.04), whereas treated tumors shifted toward *myeloid-centric* states (OR = 2.34, p = 0.01), consistent with lymphocyte depletion and myeloid expansion under therapeutic pressure. No major differences in TME distribution were observed by sex (p = 0.94) or between adult and pediatric tumors (p = 0.42, Supplementary Fig. 15a-d). Together, these data show that TME composition varies by tumor type, treatment, and disease stage, with implications for immunotherapy.

All therapeutic clusters were represented across the five major TME archetypes, reflecting substantial diversity in immune and stromal composition (Fig. 5c; Supplementary Fig. 16a-b). *Immune-stromal* TMEs were preferentially enriched in TC1 and TC4, sharing predicted drug sensitivity to valrubicin, afatinib, tanespimycin, AZD5363, and birinapant, whereas immune-rich TMEs were more prevalent in TC2, TC3 and TC5, uniquely sensitive to onvansertib (NMS-1286937) (Fig. 2b). As mentioned before, TC1-3 exhibited sensitivity to chromatin-modifying agents, including HDAC inhibitors such as romidepsin and panobinostat, consistent with their capacity to reprogram fibroblast-rich immunosuppressive TMEs (TC1)^46^ and enhance tumor immunogenicity through increased antigen presentation^50^. TC5 was markedly enriched in the *myeloid-centric* archetype, indicative of a myeloid-driven immunosuppressive TME associated with TC5 subclones. TC9 was uniquely enriched in T-cell-centric TMEs, particularly in KIRC (Supplementary Fig. 14).

To gain higher resolution, deeper subclassification with the novel TME archetypes defined in TCCA revealed additional heterogeneity within these broad TME groups. At this finer-grained level, *myeloid-centric* with Mp and Mo bias archetypes were specially enriched in TCs 3 and 5, while TC4 was dominated by *CAF-like immune-stromal* phenotype. In contrast, *immune-stromal desert* TMEs were most frequent in TC9-10 (Supplementary Fig. 16b). Analysis of sample distribution revealed that all TME archetypes were represented regardless of the degree of therapeutic heterogeneity (Supplementary Fig. 17a). Tumors spanning 2-3 TCs showed a modest increase in *immune-rich* and *T-cell-centric* TMEs compared with those confined to a single TC (n = 205 samples), which displayed diverse TMEs, suggesting that immune-infiltrated TMEs may reflect therapeutic subclonal diversity (Supplementary Fig. 17b).

Predicted immunotherapy (ICI) sensitivity was assessed with Beyondcell using the Wang et al. signature^51^, which distinguishes immunologically “hot” versus “cold” tumors and is higher in ICI responders. *Immune-desert* and *immune-stromal* tumors showed the lowest predicted ICI responsiveness, whereas T-cell-centric tumors showed the highest, revealing TME subtype-specific differences in potential immunotherapy responsiveness (Fig. 5d). Interestingly, *immune-rich*, *T-cell-centric*, and *immune-stromal* tumors exhibited higher CNA-based subclonal diversity than myeloid-centric or immune-desert tumors (Supplementary Fig. 4e), consistent with prior reports linking immune-active TMEs to genomic heterogeneity^52^. These findings indicate that therapeutic vulnerabilities are primarily encoded in malignant transcriptional programs, with TME acting as a modulatory rather than deterministic factor.

### Therapeutic clusters stratify patient outcomes and reveal a proliferative, high-risk transcriptional state

To evaluate the clinical relevance of scTherapy-derived therapeutic clusters, we examined the expression of their marker genes and assessed their enrichment in TCGA samples using GSVA and analyzed the correlation between cluster signatures and patient survival. Our results revealed that TC4, TC5, and TC10 gene signatures were significantly associated with worse progression-free interval (PFI), in contrast TC6 and TC8, which correlated with better prognosis (Fig. 6a). Notably, TC10 exhibited the strongest association with poor clinical outcomes. When analyzed by tumor type, TC10’s signature was linked to worse prognosis in the majority of cases, with this trend being statistically significant in 10 of 30 cancer types (Supplementary Fig. 18).

**Figure 6.**
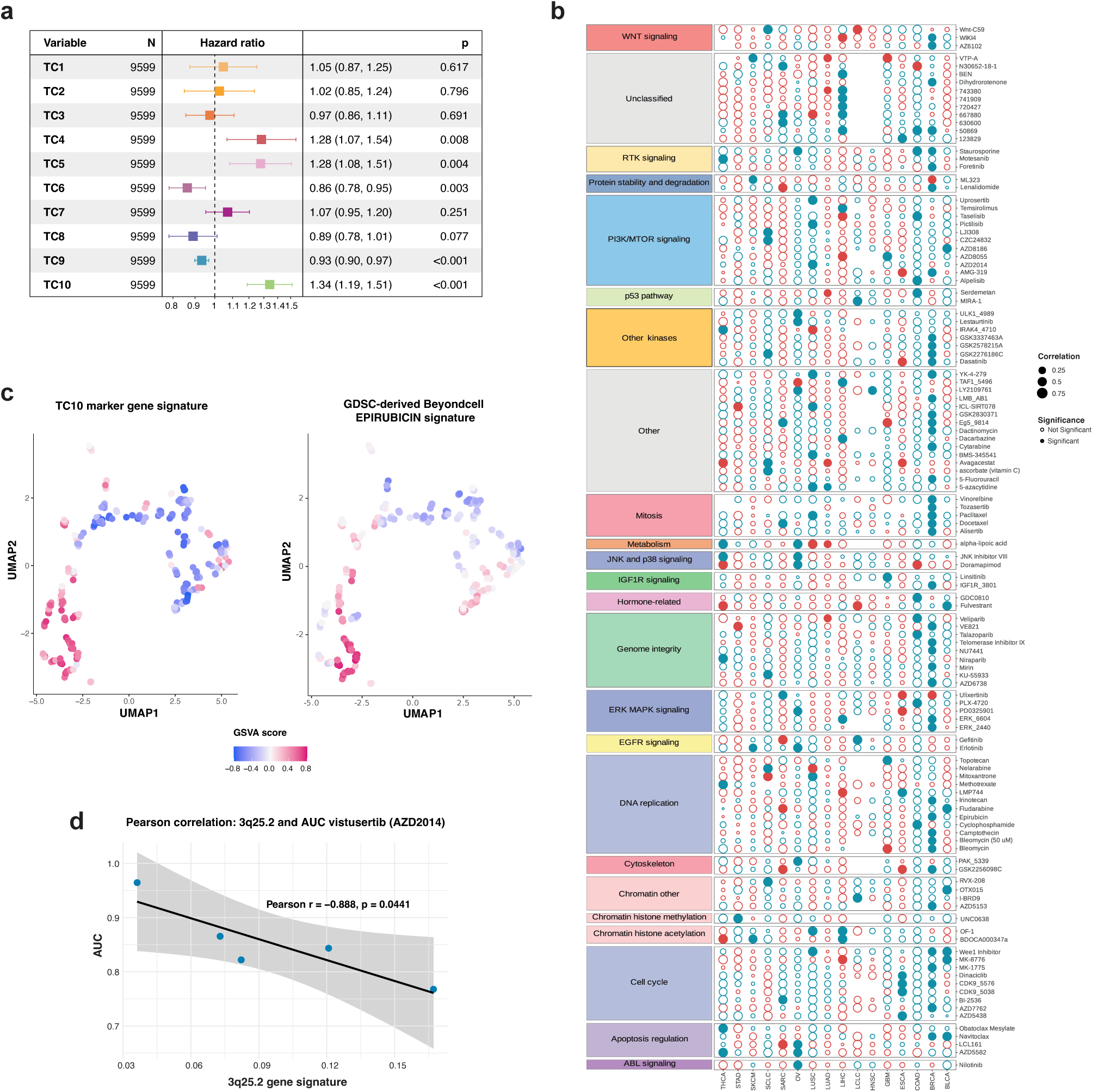
Integration of genomic, transcriptomic, and TME features with drug response in therapeutic clusters TC4 and TC10. **a** Forest plot showing the survival risk associated with the expression of TC1–TC10 marker genes in TCGA tumors (Cox proportional hazards models adjusted for age, sex, cancer type, clinical stage, and tumor grade). **b** Correlation between TC10 marker expression and drug response in cancer cell lines from the GDSC dataset. The y-axis shows cancer types; the x-axis shows GDSC drugs grouped and color-coded by mechanism of action. Only drugs showing at least one significant negative correlation with TC10 marker expression in the corresponding cancer type are displayed (i.e., higher TC10 signature expression is associated with lower AUC values, indicating greater drug sensitivity). **c** Signature enrichment in subclone pseudobulks of BRCA patient cells (n = 258 subclones). Left: UMAP showing enrichment of the TC10 marker gene signature. Right: UMAP displaying GSVA scores of the GDSC-derived Epirubicin signature from the Beyondcell drug sensitivity collection (SSc). **d** Pearson correlation between the expression of band 3q25.2 and response to AZD2015 (or vistusertib, a PI3K/mTOR inhibitor predicted by scTherapy) across five ESCA cell lines from Kinker et al^6^.

TC10 is the smallest group (n = 101 subclones), comprising a mix of cancer types with high transcriptomic heterogeneity but remarkably homogeneous therapeutic profiles (Fig. 2e). The most frequently predicted drug mechanisms of action for this cluster include inhibitors of the ubiquitin-proteasome system (carfilzomib), DNA-damaging agents (epirubicin, mitoxantrone), PI3K/AKT/mTOR pathway inhibitors, and a kinase inhibitor (Fig. 2b). This cluster is characterized by a highly proliferative phenotype marked by the expression of genes like MCM7, PCNA and CDK4 (Fig. 2c). As expected from its relatively low genomic heterogeneity (Fig. 2e), recurrent CNV events are rare, with none detected in over 40% of subclones.

To investigate TC10 therapeutic vulnerabilities, we integrated Genomics of Drug Sensitivity in Cancer (GDSC) drug sensitivity data with TC gene signature enrichment across cancer cell lines using the Kinker et al. single-cell dataset. This identified 142 significant drug-tumor type associations, with BRCA cell lines showing the highest number of significant drug correlations (Fig. 6b). Many drugs targeted pathways predicted by scTherapy as relevant for TC10, including cell cycle regulation, DNA replication, and genome integrity maintenance. Epirubicin, a topoisomerase II inhibitor, showed a strong inverse correlation with TC10 signature enrichment in BRCA lines. Epirubicin, an anthracycline widely used in breast cancer chemotherapy particularly in the adjuvant setting and in patients with metastatic disease, has demonstrated efficacy against highly proliferative tumors^53^. This suggests that it may be particularly effective in more advanced or aggressive stages of disease, consistent with the poor prognosis associated with TC10 and the EMT-related pathways enriched in this cluster. Applying Beyondcell to BRCA subclones (pseudobulk) revealed that subclones with higher TC10 marker expression exhibited increased sensitivity to epirubicin (Fig. 6c), validating scTherapy predictions. Further analysis of TC10 distribution in breast cancer revealed that TC10 subclones were almost exclusively found in the triple-negative breast cancer (TNBC) subtype, known for its aggressive nature and high proliferative capacity (Supplementary Fig. 19). These results identify TC10 as an aggressive, clinically relevant cell population across cancer types, particularly TNBC, whose sensitivity to cell cycle-targeting therapies highlights potential avenues for targeted treatment.

### 3q-driven vulnerabilities define a therapeutically homogeneous and actionable subtype of esophageal cancer

TC4, enriched in esophageal cancer (ESCA) subclones, emerged as a therapeutically homogeneous yet biologically distinctive cluster, enriched in stress-related and plasticity-associated pathways (Fig. 2a, Fig. 4c). TC4 subclones share common drug sensitivities, including those to chromatin-modifying and cell cycle arrest agents and kinase inhibitors such as abemaciclib and ceritinib, an ALK inhibitor consistent with the presence of ALK amplifications in ∼11% of esophageal cancers^54^ (Fig. 2b). Predicted sensitivities also included PI3K/AKT/mTOR pathway inhibitors (AZD2014, MLN0128, PI-103) and EGFR inhibitors like afatinib, shared with TC1, reflecting convergent oncogenic signaling dependencies. At the genomic level, TC4 was characterized by recurrent amplifications along chromosome 3q, specifically at 3q25.1-3q29 and deletions on 5q, alterations commonly observed in esophageal squamous cell carcinomas^55^ (Fig. 2d, Supplementary Fig. 20a). Two subclonal amplification patterns of the 3q25.1-3q29 region were also identified: one lacking 3q25.1-3q25.2, and another restricted to these regions (Supplementary Fig. 20b). Survival analysis in TCGA ESCA samples revealed that high expression of genes within these 3q regions correlated with poorer prognosis, underscoring their role in EMT and malignant progression (Supplementary Fig. 21). Analysis of ESCA cell lines from the GDSC database showed that amplifications spanning 3q25.1-3q26.33 were associated with increased sensitivity to apoptosis inducers (venetoclax), cell cycle inhibitors (dinaciclib) and chromatin regulators (GSK-LSD1) (Supplementary Fig. 22). Importantly, although the overlap between GDSC-tested compounds and scTherapy predictions was limited, the PI3K/mTOR inhibitor vistusertib (AZD2014) was consistently identified by both datasets, providing cross-validation of this therapeutic dependency in TC4 (Fig. 6d). Intersecting these cytobands with TC4 marker genes (Fig. 2c) identified several potential mediators of drug response. *RAP2B* (3q25.2), RAS-family GTPase and p53 target gene, regulates cytoskeletal remodeling and confers pro-survival function; consequently, targeting RAP2B may sensitize tumor cells to DNA damage-induced apoptosis and PI3K/mTOR pathway inhibitors^56,57^ (Supplementary Fig. 22a). *TM4SF1* and *PFN2* (3q25.1) promote cell motility and EMT through activation of the PI3K/AKT/mTOR pathway, providing a mechanistic basis for mTOR inhibitor sensitivity in tumors with 3q amplification^58,59^. *ACTL6A* (3q26.33), a core subunit of the SWI/SNF chromatin remodeling complex frequently amplified in esophageal squamous cell carcinoma, may confer sensitivity to chromatin-modifying agents^60^. Additional genes within 3q27.1-3q29, including *PSMD2* (proteasome activity) and *MAP3K13* (JNK/p38 signaling), further implicate cell cycle, stress, and proteostasis pathways in TC4 drug response. Amplification of 3q27.1 also correlated with reduced EGFR inhibitor sensitivity (Supplementary Fig. 22b), suggesting potential cross-resistance mechanisms^61^.

TC4 is marked by a highly stromal composition, with a dominant CAF-rich TME, coupled with immune components such as cDC1, cDC2, and Tregs, indicating stromal remodeling and immune suppression that may constrain therapeutic efficacy (Fig. 5c; Supplementary Fig. 16a-b). TC4 expressed basal and squamous epithelial markers (*KRT5*, *KRT6A/B*, *DSG3*, *PKP1*), defining a keratinized, basal-like phenotype with partial EMT features characteristic of esophageal squamous epithelium^62^ (Fig. 2c). Together, these results position TC4 as a biologically coherent and therapeutically actionable subgroup, where convergent genomic amplifications drive coordinated transcriptional states and drug vulnerabilities. Despite its fibrotic and immunosuppressed TME, the combination of chromatin, PI3K/mTOR, and cell cycle inhibitors may represent a rational therapeutic strategy for esophageal tumors harboring 3q amplifications, with potential implications for other cancers harboring similar genomic alterations.

## Discussion

Understanding how ITH influences therapeutic response remains a central challenge in oncology^63,64^. Accumulating evidence indicates that therapy response emerges from interactions between genetic diversity, transcriptional plasticity, and the TME on both malignant and non-malignant compartments rather than genomic alterations alone^65–69^. However, a systematic, pan-cancer framework linking these dimensions to drug response at single-cell resolution has been lacking.

Here, we introduce the Therapeutic Cancer Cell Atlas (TCCA) as a unified single-cell resource to map therapeutic heterogeneity across cancers. By integrating clonal architecture, transcriptional programs, inferred drug sensitivities, and TME archetypes across ∼1.8 million cells, we provide a multidimensional view of therapeutic vulnerabilities across cancers. A key finding of this study is that therapeutic heterogeneity is largely decoupled from genomic and transcriptomic heterogeneity, instead arising from recurrent functional programs and cellular contexts. Tumors with a moderate number of subclones such as BRCA and LUAD can exhibit broad drug-response divergence, while genomically heterogeneous tumors may converge on shared vulnerabilities^66^.

A major contribution of TCCA is the identification of ten recurrent therapeutic clusters that capture conserved and context-specific drug dependencies across tumor types. These clusters reveal shared vulnerabilities that transcend tissue of origin, alongside lineage-restricted therapeutic states. For example, TC1-3 define epithelial-associated clusters with overlapping sensitivities, supported by shared transcriptional programs and recurrent amplifications. In contrast, TC5 links primary gliomas and melanoma-derived brain metastases through convergent dependencies on cell-cycle and PI3K signaling, illustrating how anatomical niche and functional state can override tissue origin in shaping therapeutic response. Importantly, TCCA also identifies clusters characterized by high therapeutic homogeneity, such as TC4 and TC10, which may represent particularly tractable targets for precision treatment. TC4 delineates a biologically coherent subtype of esophageal cancer driven by 3q amplifications and associated with coordinated sensitivity to chromatin, PI3K/mTOR, and cell-cycle inhibitors. TC10, despite comprising subclones from multiple cancer types, exhibits a shared proliferative transcriptional state, poor clinical outcomes, and consistent sensitivity to DNA-damaging and proteasome-targeting agents. The strong association of TC10 with adverse prognosis across cancers underscores the clinical relevance of functional therapeutic states that cut across histological boundaries. Conversely, clusters with high therapeutic heterogeneity, particularly TC6-TC9, highlight challenges for treatment design. These clusters harbor subclones with divergent drug sensitivities driven by stress-response and proteostasis programs, underscoring the need for combination or adaptive treatment strategies. Together, these findings illustrate that therapeutic heterogeneity is structured and can be organized into recurrent functional states with predictable drug-response patterns.

By integrating transcriptional metaprograms, we further show that therapeutic clusters are underpinned by conserved programs of malignant plasticity. Programs related to proliferation, MYC activation, EMT, inflammation, stress, and lineage identity systematically align with drug sensitivities, providing mechanistic insight into why specific clusters respond to particular therapies. These results reinforce the notion that functional transcriptional states act as intermediaries between genotype and drug response, offering a more actionable framework than histology or mutation-based stratification alone.

The TME emerged as an additional determinant of therapeutic diversity. The TCCA recapitulates known TME archetypes while defining novel immune and stromal subtypes that further refine therapeutic behavior. Although all therapeutic clusters span multiple TME states, certain associations emerge for example, CAF-rich immune-stromal TMEs in TC4 and TC1, or myeloid-centric TMEs in TC5. Notably, therapeutic vulnerabilities persist across diverse TMEs. These results suggest that malignant transcriptional programs play a dominant role, while the TME modulates accessibility, immune engagement, and resistance. Therapy and metastatic progression were associated with immunosuppressive TME shifts, underscoring the need for context-aware combination strategies. Pediatric tumors are enriched for proliferative, tumor-specific clusters, whereas adult tumors display more diverse landscapes. Age, sex, and metastatic site showed limited impact, emphasizing the primacy of cellular and microenvironmental states in shaping drug response.

The clinical relevance of TCCA is underscored by its ability to stratify patient outcomes and nominate actionable therapies. Linking TC signatures to TCGA survival data identifies high-risk transcriptional states, while integration with pharmacogenomic datasets and single-cell drug sensitivity models validates predicted vulnerabilities. The epirubicin-TC10-TNBC example illustrates how TCCA can bridge single-cell inference and clinically used agents, highlighting opportunities for drug repurposing and rational patient stratification. More broadly, the atlas enables patient-level dissection of therapeutic heterogeneity, revealing how genomic complexity, subclonal composition, and functional state jointly influence drug responsiveness in both primary and metastatic disease. Despite these insights, translating TCCA predictions into clinical benefit requires addressing key methodological limitations and validating functional hypotheses. Predictions rely on transcriptomic inference and may not capture metabolic or post-translational determinants of drug efficacy. Cross-sectional datasets limit assessment of temporal dynamics and resistance evolution. Future studies incorporating longitudinal sampling, perturbation experiments, and patient-derived functional assays will be essential to refine cluster-specific vulnerabilities and validate combination strategies. Nonetheless, the strong concordance between inferred sensitivities, pharmacogenomic datasets, and clinical associations supports the robustness of the TCCA framework.

In summary, the TCCA provides a comprehensive, single-cell map of therapeutic heterogeneity across cancers, linking subclonal functional states, transcriptional programs, and microenvironmental context to drug response. This resource enables tumor- and drug-centric exploration of vulnerabilities, identifies resistant compartments, and supports rational design of adaptive and combinatorial therapies. The full dataset and interactive resource are publicly available (https://tcca.bioinfo.cnio.es/), facilitating community-driven exploration and accelerating the translation of single-cell insights into clinical impact.

## Methods

### Data collection and inclusion criteria

For data selection, we conducted a search on PubMed for scRNA-seq datasets that provided publicly available data (for example, through the Gene Expression Omnibus database repository). All selected publications must freely provide the data in the form of an expression matrix, together with a list of associated genes and cells (barcodes). In addition, we prioritized datasets that included multiple patient tumours, with annotated cell types and a substantial proportion of malignant cells. A total of 36 studies meeting these criteria were identified, comprising 1820 tumor samples and approximately ∼5,5 million single cells, which were subsequently preprocessed. For each sample, we extracted available metadata from the original publications, including the sample and patient identifiers, cell type annotations, and malignancy status, when available. Finally, to ensure robustness, we retained only samples containing at least 100 malignant cells and with available clinical annotations. This filtering step resulted in 853 samples comprising approximately 1.8 million cells, including 1 million malignant cells and 800,000 tumor microenvironment (TME) cells.

### Clinical database construction

A comprehensive clinical database was constructed by manually curating sample-specific clinical information from 36 original research papers. An expert clinician extracted and standardized metadata. This included demographic data (sample ID, study, patient, sex, age), tumor characteristics (type, subtype, histopathological grade, TNM classification, anatomical site, primary vs. metastatic status), treatment history (prior therapy, treatment type, response), and survival metrics (overall survival and progression-free interval), when available (Supplementary table 1). To enable cross-study comparisons, the 46 initial tumor types were consolidated into 34 broader “refined tumor type” categories using TCGA nomenclature when applicable (e.g., astrocytomas and oligodendrogliomas grouped as LGG). Technical metadata, including sequencing platform and reference genome, were also recorded to facilitate downstream scRNA-seq analyses.

### Preprocessing of scRNA-seq data

Each study was preprocessed independently using Seurat (v4.4.0) within a Snakemake pipeline (v8.23.0). Cells were excluded if they exhibited mitochondrial content >10%, ribosomal content >40%, <1,000 or >7,000 expressed genes, or <500 total counts. Genes expressed in <5% of cells were filtered out. Filtered data were log-normalized (NormalizeData), variable features identified (FindVariableFeatures), and expression matrices scaled (ScaleData) using default parameters. Clinical metadata, including sample/patient identifiers, cell type annotations, and malignancy status, were integrated into each Seurat object.

To optimize large-scale analysis, we migrated to Seurat v5.3.0 with BPCells v0.2.0, enabling in-memory sketching for rapid analysis while maintaining full datasets on-disk. Samples with <100 malignant cells or missing clinical annotations were excluded. For samples lacking sex annotation (n=203), we inferred sex using the classifySex function from Speckle (v1.2.0), a logistic regression classifier trained on mouse and single-cell RNA-seq data. Sample-level sex was assigned when ≥80% of cells showed concordant predictions, successfully annotating 151 samples.

### Inference of Copy Number Alterations (CNAs) and malignant cells annotation

To assess the clonal structure of the samples in the cohort, we used SCEVAN (v1.0.1). The pipeline was run independently for each sample to infer their copy number alteration (CNA) profiles across all autosomes, as sex chromosomes were not considered by SCEVAN. As a result, CNAs were successfully analyzed in 95% of the samples; the remaining 5% failed due to unrecoverable errors. Subclone and malignancy annotations provided by SCEVAN were incorporated into the cohort metadata and used in downstream analyses.

In addition, the pipeline generated both cell-by-gene and subclone-by-region matrices for each sample. The resulting subclone-by-region matrices were harmonized by converting the genomic regions reported by SCEVAN into cytobands using the GenomicRanges Bioconductor package. Specifically, SCEVAN-inferred genomic segments were mapped to cytogenetic bands (cytobands) from the hg38 reference genome using positional information retrieved through AnnotationHub. Each segment was intersected with overlapping cytobands using the join_overlap_intersect function from the plyranges package. When multiple segments mapped to the same cytoband, the CNA values were averaged across overlapping segments.

### Cell type annotation

We performed automated cell type annotation using Azimuth (v0.5.0), which maps cells to reference datasets at two resolution levels: broad (main cell type) and refined (fine cell type). Annotation confidence was assessed using Azimuth’s mapping score (how well cells align to the reference) and prediction score (confidence in assigned labels). Cells with mapping scores ≥0.7 were considered high-confidence annotations. For cells with multiple high-confidence matches across references, we selected the annotation with the highest prediction score. Cells below the 0.7 threshold retained the best-available low-confidence annotation.

We then harmonized both author-provided and Azimuth-inferred annotations into a unified nomenclature using manually curated reference tables. Author annotations were prioritized when available; otherwise, Azimuth annotations were used. Of 62,615 cells lacking author annotations, 18,251 also lacked high-confidence Azimuth mapping and were annotated using low-confidence predictions. For two datasets where Azimuth performed particularly poorly (pdac_shu_zhang and skcm_chao_zhang), we instead re-annotated all cells using SingleR (v2.0.0) and harmonized labels to the common scheme.

Since Azimuth references derive from healthy tissues and lack malignant cells, we excluded malignant populations from automated annotation. These cells were manually labeled as “Malignant” by harmonizing author-specific labels (e.g., epithelial basal/luminal cells, RCC) or, when unavailable, using SCEVAN predictions based on copy number alterations. The final annotation comprised 21 main cell types and 140 refined subtypes, validated by examining marker gene expression in each population.

### Dimensional reduction and integration

We transitioned to Python-based tools for dimensionality reduction and clustering due to their efficiency with large-scale datasets. The raw expression matrix and metadata from 1.8 million cells were extracted from Seurat and compiled into an h5ad object (tcca_expression_metadata_lvl2.h5ad). Using Scanpy (v1.10.1), we log-normalized the data, identified 2,000 highly variable genes (Seurat v3 method), performed principal component analysis (PCA), constructed a nearest-neighbor graph, and generated UMAP embeddings.

Initial visualization revealed substantial batch effects across samples from different studies. To address this, we benchmarked three integration methods: scVI and scANVI (scvi-tools v1.1.2), both deep learning approaches, and Scanorama (v1.7.4), a linear embedding method. The scVI model was trained with two hidden layers and 30-dimensional latent space, using a negative binomial likelihood over 50 epochs. The scANVI model, which incorporates cell type labels during integration, was similarly trained for 50 epochs. Additionally, Scanorama was applied with default parameters.

Integration performance was evaluated using scIB (v0.5.0), which quantifies both biological conservation and batch correction. Based on these metrics and prior benchmarking studies, we selected scANVI as it provided the best balance of batch correction and biological signal preservation. Post-integration, cells from the same type clustered appropriately, confirming effective batch correction. All analyses were performed on a High Performance Cluster (HPC) with 3× Nvidia A100 80GB GPUs.

### Drug sensitivity prediction

We applied two complementary methods to predict drug response in malignant cells.

#### Beyondcell-based prediction

We used Beyondcell (v2.1.0) to compute drug sensitivity scores using single-cell expression data with a collection of drug sensitivity signatures (SSc) covering 581 drugs. Due to scalability constraints, Beyondcell was run separately on each of the 36 studies via a Snakemake pipeline. Individual sensitivity score matrices were merged into a comprehensive BPCell matrix (581 drugs × ∼1 million malignant cells), converted to a Seurat object, and analyzed using a representative sketch of 50,000 cells. Clustering was performed across multiple k and resolution values, with performance evaluated using modularity, silhouette score, Davies-Bouldin index, and clustering purity. Optimal parameters were k = 300 neighbors and resolution = 0.5. To enable comparison with scTherapy, we also computed subclone-level pseudobulk profiles and applied Gene Set Variation Analysis (GSVA) with the SSc signature collection.

#### scTherapy-based prediction

For primary downstream analysis, we selected scTherapy to reduce complexity arising from high intratumoral heterogeneity. scTherapy generates drug response predictions at the sample-specific subclone level, providing a more interpretable framework. We included only cells consistently labeled as malignant by both original authors and SCEVAN (n = 637,062 cells). Study-specific Seurat objects were split by sample, retaining only samples with both malignant and healthy cells. Within each sample, cells were grouped into subclones using SCEVAN assignments. As part of the scTherapy workflow, differential expression analysis was performed between each subclone and matched healthy cells using FindMarkers (Wilcoxon rank-sum test). DEGs were filtered using two criteria: (i) adjusted p-value ≤ 0.05 with |log2FC| > 1 for significantly altered genes, and (ii) log2FC between –0.1 and 0.1 for stably expressed genes. Drug predictions for each subclone were generated using scTherapy’s predict_drugs() function with exclude_low_confidence = TRUE, which applies a pre-trained LightGBM model built on large perturbation datasets. All predictions were aggregated into a global table (sctherapy_drug_predictions_subclone.tsv) spanning 2,627 subclones across all samples.

### Generation of therapeutic clusters and identification of top candidate drugs

To identify groups of subclones with similar predicted drug response profiles, pairwise Jaccard indices were computed between their predicted drug sets. Spectral clustering (specc, kernlab v0.9-32) was then applied to the resulting similarity matrix. After testing multiple cluster numbers, ten clusters were selected to balance biological interpretability with sufficient granularity for therapeutic stratification.

For each drug response cluster, marker genes were identified by differential expression analysis using FindAllMarkers (Wilcoxon rank-sum test) comparing subclonal cells within a cluster against all other subclones. Only positive markers were retained and filtered using the following criteria: avg_log2FC ≥ log2(1.5), pct.1 ≥ 0.5, and pct.1 - pct.2 ≥ 0.20.

Representative drugs for each cluster were selected as those predicted in >60% of subclones within the cluster. The top drugs per cluster were then annotated with their known mechanisms of action (MoAs).

### Transcriptomic, genomic, and therapeutic ITH quantification

We quantified intratumoral heterogeneity (ITH) at three levels—transcriptomic, genomic, and therapeutic—using a consistent two-step framework: first computing subclone-level variability, then aggregating to sample-level ITH weighted by subclone cellular frequency.

#### Transcriptional ITH (tITH)

Transcriptional heterogeneity was quantified using the Coefficient of Variation (CV) of highly variable genes (HVGs). First, 3,000 HVGs were identified across the full dataset using a variance-stabilizing transformation (FindVariableFeatures, Seurat). To ensure robustness, only genes with a mean log-normalized expression >0.01 across all cells within each subclone were retained. For each subclone *k*, gene-level CVs were computed from log-normalized expression values across all cells belonging to that subclone (*n*_*k*_ ≤ 3), and the subclone-level transcriptional heterogeneity 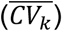 was defined as the mean CV across all retained HVGs. Sample-level tITH was calculated as the weighted mean of subclone CVs, where weights correspond to cellular frequencies (*f*_*k*_):

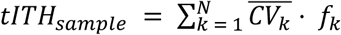

where N is the total number of subclones in the sample, 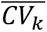 is the mean CV of subclone *k*, and *f*_*k*_ denotes the relative cellular frequency of subclone *k*, defined as the number of cells in subclone *k* (*n*_*k*_) divided by the total number of cells in the sample (*n*), such that *f*_*k*_ = *n*_*k*_/*n*.

#### Genomic ITH (gITH)

Genomic ITH was quantified using Shannon Entropy derived from copy number variation (CNV) profiles across 790 genomic cytobands (chromosomes 1–22). For each subclone, raw CNV values were discretized into 5 equal-width bins, and subclone-level entropy (*H*_*k*_) was computed as:

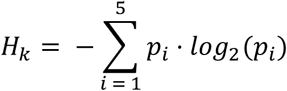

where *p*_*i*_ is the proportion of the subclone’s cytobands in bin *i*. Sample-level gITH was calculated as the Shannon entropy of the weighted CNV distribution, where each bin’s proportion was weighted by subclone cellular frequencies (*f_k_*): *Weighted prop*_*i*_ = ∑ _*k*∈ *sample*_(*p*_*k*,*i*⋅_ · *f*_*k*_) where *p*_*k*,*i*_ represents the same proportion *p*_*i*_ calculated specifically for subclone k. The final sample-level gITH was computed as the Shannon Entropy of this resulting weighted distribution:

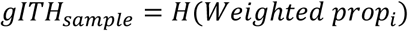

This approach measures the overall genomic disorder of the tumor by accounting for the complexity of individual subclone CNV profiles and their relative abundance.

#### Therapeutic ITH (thITH)

Therapeutic heterogeneity was quantified based on dissimilarity of drug response predictions between subclones. For each subclone, intrinsic therapeutic variability was measured as the mean Jaccard distance between its drug response profile and all other subclones within its assigned therapeutic cluster. Sample-level thITH was computed using Rao’s Quadratic Entropy (Q), which accounts for both pairwise therapeutic dissimilarity (Jaccard distance matrix D) and cellular frequencies (*f*_*k*_):

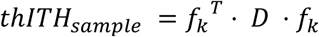

This metric provides a single measure of the overall therapeutic diversity within the sample, weighted by subclone abundance.

### Identifying recurrent functional gene programs across tumour samples

We investigated transcriptional programs of intratumoral heterogeneity in the malignant cells+ using non-negative matrix factorization (NMF). For each tumour sample, the UMI counts were normalized to CPM (counts per million). The CPM matrix was then log2-transformed, as E_i,j_=log2(CPM_i,j_/10+1), for gene i in cell j, and filtered to retain genes with E_i,j_ > 3.5 in at least 2% of cells. Expression values were then mean-centered and negative values were set to zero before applying NMF (RcppML v.0.3.7) across multiple factorization ranks (K = 4–9), generating 39 programs for each tumor sample. Each program was summarized by its top 50 genes based on the NMF gene weight matrix. Following Kinker et al. approach for identifying recurrent heterogeneous programs (RHPs) of gene expression^6^, we defined robust NMF programs as those that are reproducible (≥70% gene overlap across NMF ranks), non-redundant (filtered for ≥20% overlap within samples), and recurrent (≥20% overlap across samples). This approach yielded a total of 4585 robust NMF programs.

To identify metaprograms (MPs), we clustered robust NMF programs by gene overlap using the custom approach from Gavish et al^8^, generating 66 initial MPs. After removing MPs enriched for ribosomal/mitochondrial genes, those with ≤10 programs, MPs derived from a single study when that cancer type was represented in multiple studies, or those resembling non-malignant cell types, we retained 43 MPs (Supplementary table 2). These were functionally annotated using overrepresentation analysis (ORA) with MSigDB collections (H, C2, C4, C5.GOBP, C5.GOCC, C5.GOMF, C8) and published gene signatures from cancer cell states^5–8^. ORA was performed using the fora function from the fgsea package (v1.24.0), with gene sets considered significantly enriched at FDR-adjusted P < 0.05 (hypergeometric test; Supplementary table 3). The 43 MPs were subsequently grouped into 13 families based on functional similarity.

MP enrichment in malignant cells was quantified using AddModuleScore_UCell. For each therapeutic cluster, we calculated mean UCell scores, the percentage of cells exceeding the MP-specific 75th percentile, and z-scores standardized and rescaled to 0–1. We also evaluated CIN70 signature enrichment to estimate chromosomal instability per therapeutic cluster.

### Sample-specific tumor microenvironment classification

Following the approach of Combes et al., we classified samples into immune archetypes based on TME composition. While the original method used gene signatures to infer cell-type proportions from bulk RNA-seq data, we directly quantified immune and stromal cell-type fractions from scRNA-seq annotations. Liquid tumors (LAML, ALL, MM, CLL) and samples lacking non-malignant cells were excluded, yielding 493 solid tumor samples for TME classification.

For each sample, we computed the relative fraction of cells belonging to 12 consolidated TME cell-type categories: Lymphoid, CD8T, CD4T, Treg, Stromal, CAF, Endothelial, Myeloid, Macrophage (Mp), Monocyte (Mo), cDC1, and cDC2. This generated a 493 samples × 12 TME features matrix, which was used to construct a Seurat object. We performed PCA on scaled data and built a k-nearest neighbor (KNN) graph using FindNeighbours (Euclidean distance in PCA space, k = 3–300). Clustering was performed using the Louvain algorithm (FindClusters) across resolution parameters from 0.3 to 2.0 (0.1 increments).For each k and resolution combination, we evaluated clustering quality using the Davies-Bouldin Index (DBI), which quantifies cluster separation by the ratio of intra- to inter-cluster distances. The optimal clustering (lowest DBI) was achieved at k = 7 and resolution = 0.7, yielding 12 robust TME clusters. Clusters were manually annotated based on dominant TME profiles, resulting in biologically interpretable archetypes: *“immune rich”, “immune rich Treg, cDC2 bias”, “T-cell centric”, “myeloid centric”, “myeloid centric Mp bias”, “myeloid centric Mo bias (high)”, “myeloid centric Mo bias (medium)”, “immune stromal”, “immune stromal endo-like”, “immune stromal CAF-like Mp bias”, “immune desert CAF-like”, and “immune stromal desert”*. Each sample was assigned to a single TME archetype based on its cluster membership.

### Correlation of therapeutic clusters with survival

To assess the prognostic value of therapeutic clusters, we leveraged bulk RNA-seq data from TCGA pan-cancer cohorts. Gene expression data (HTSeq counts) were normalized using TMM normalization (edgeR package) and voom transformation (limma package). For each therapeutic cluster, we computed patient-level enrichment scores using single-sample Gene Set Enrichment Analysis (ssGSEA) via the GSVA package with cluster-specific marker gene signatures. Cox proportional hazards models were fitted to evaluate associations between signature enrichment and survival outcomes (overall survival or progression-free interval, depending on cancer type), adjusting for cancer type, sex, age, tumor stage, and histological grade. For cancer-specific analyses, models were stratified by tumor type. Kaplan-Meier curves were generated by dichotomizing patients into high and low signature groups based on optimal cutpoints determined using maximally selected rank statistics (survminer package).

### Validation of therapeutic predictions using experimental data from GDSC

To validate therapeutic cluster sensitivities, we selected the 102 cell lines common between the dataset used in Kinker et al. 2020 and the Genomics of Drug Sensitivity in Cancer (GDSC) database. For these cell lines, we used the single-cell data to calculate the enrichment scores of each cluster’s biomarker signature at the single-cell level using the UCell package. Subsequently, we computed the average enrichment per cell line and calculated the Pearson correlation between the resulting enrichment values and the drug area under the curve (AUC) values from GDSC. Correlations were calculated within each cancer type by grouping the cell lines accordingly. TC10-specific drugs were defined as those showing a negative correlation between the TC10 signature and AUC, but not showing this correlation with any other TC signature.

The same approach was applied to gene signatures derived from selected cytobands associated with TC4. These signatures were generated by retrieving all genes located within specified chromosomal cytobands using the Ensembl database via biomaRt. In this case, correlations with AUC values were specifically computed for esophageal cancer cell lines.

### Statistical analysis

All statistical tests, significance thresholds, and exact n values are reported in figure legends, results, or methods sections. Sample inclusion was based on data availability (publicly accessible scRNA-seq datasets with clinical annotations and ≥100 malignant cells per sample); no samples were excluded except for technical filtering steps described in Methods. Clustering performance was evaluated using Davies-Bouldin Index, silhouette score, and clustering purity, with optimal parameters selected based on lowest DBI or best combined metrics when applicable. Differential expression analyses used the two-tailed Wilcoxon rank-sum test. Cox proportional hazards models were fitted to assess survival associations, with hazard ratios, 95% confidence intervals, and Wald test p-values reported. Multiple testing correction was performed using the Benjamini-Hochberg method (FDR).

## Data availability

This work relied on curation and integrative analysis of 36 external scRNA-seq studies and did not involve generation of new primary data. All curated data, clinical annotations, and downstream analysis results are publicly available through the TCCA portal at https://tcca.bioinfo.cnio.es/, which provides interactive exploration of drug response predictions, TME archetypes, and functional metaprograms. The complete integrated dataset, including a 1.8 million cell h5ad object with clinical metadata and multi-layered analysis results, as well as individual result files for genomic inference (SCEVAN CNV profiles), functional metaprograms, TME archetypes, and drug predictions (Beyondcell and scTherapy), are available at Zenodo: https://doi.org/10.5281/zenodo.18109100. Source data for all figures are provided in Supplementary tables. Original raw sequencing data from the 36 studies can be accessed through their respective GEO accession codes listed in Supplementary table 1.

## Code availability

All analysis code, including Snakemake pipelines, R and Python scripts for data integration, TME classification, CNV inference, drug response prediction, metaprogram discovery, and figure generation, is publicly available at https://github.com/cnio-bu/tcca with detailed documentation and environment specifications.

## Declarations

### Ethics approval and consent to participate

Not applicable.

### Consent for publication

Not applicable.

### Competing interests

The authors declare that they have no competing interests.

### Funding

CNIO Bioinformatics Unit is supported by Project IMPaCT-Data (IMP/00019) funded by the Spanish State Research Agency (AEI), the Carlos III Health Institute (ISCIII), co-funded by the European Regional Development Fund (ERDF), “A way of making Europe’’; the project PID2021-124188NB-I00 and PID2024-156049OB-I00/AEI/10.13039/501100011033/ funded by the Spanish Ministry of Science and Innovation/AEI, the European Union, and the ERDF “A way of making Europe”; the iTIRONET project funded by Community of Madrid (P2022/BMD-7379) and co-financed by European Structural and Investment Fund; the PRYCO234528VALI project funded by the Scientific Foundation of the Spanish Association Against Cancer (AECC); the project code Grant HR23-00051 funded by “la Caixa’’ Foundation; the PMP22/00064 project funded by ISCIII, the European Union, and the Recovery, Transformation and Resilience Plan (PRTR) through Next GenerationEU recovery funds (MRR); the project IMPaCT Digital Platform (PMP24/00024), IMPaCT-Data (PMP22/00088) and IMPaCT-VUSCan (PMP22/0006) funded by ISCIII; and the Paradifference Foundation. MG-B. is funded by the Spanish Ministry of Science, Innovation and Universities through the State Program to Develop, Attract and Retain Talent [FPU2022-01155]. IS-P. is funded by the Spanish Ministry of Science, Innovation and Universities through the LI_10 C005/24-ED CV1 NextGen Program [BEX240034].

### Author’s contributions

MG-B. designed and implemented the computational framework and conducted the majority of analyses with guidance from LS-R. F.A. and GG-L. LS-R. performed CNV inference and in silico validation of therapeutic clusters. SG-M. coordinated scRNA-seq study recruitment and manually curation the clinical information. OL-S. performed initial annotation of TME. PG-M. assisted with annotation of specific studies. IS-P. contributed to the development of the TCCA web server. MG-B., LS-R., GG-L. and F.A. wrote the manuscript. F.A., GG-L. and LS-R. supervised the project. F.A. initiated and led the study. All authors read and approved the final manuscript.

## Acknowledgments

We thank all members of the CNIO Bioinformatics Unit for their continuous support. We are also grateful to María José Jiménez-Santos for her helpful suggestions and discussions about the state of the art.

## Notes

https://tcca.bioinfo.cnio.es

https://zenodo.org/records/18109100

https://github.com/cnio-bu/tcca.git

### Competing Interest Statement

The authors have declared no competing interest.

