## Supplementary Figures for "A single-cell atlas linking intratumoral states to therapeutic vulnerabilities across cancers"

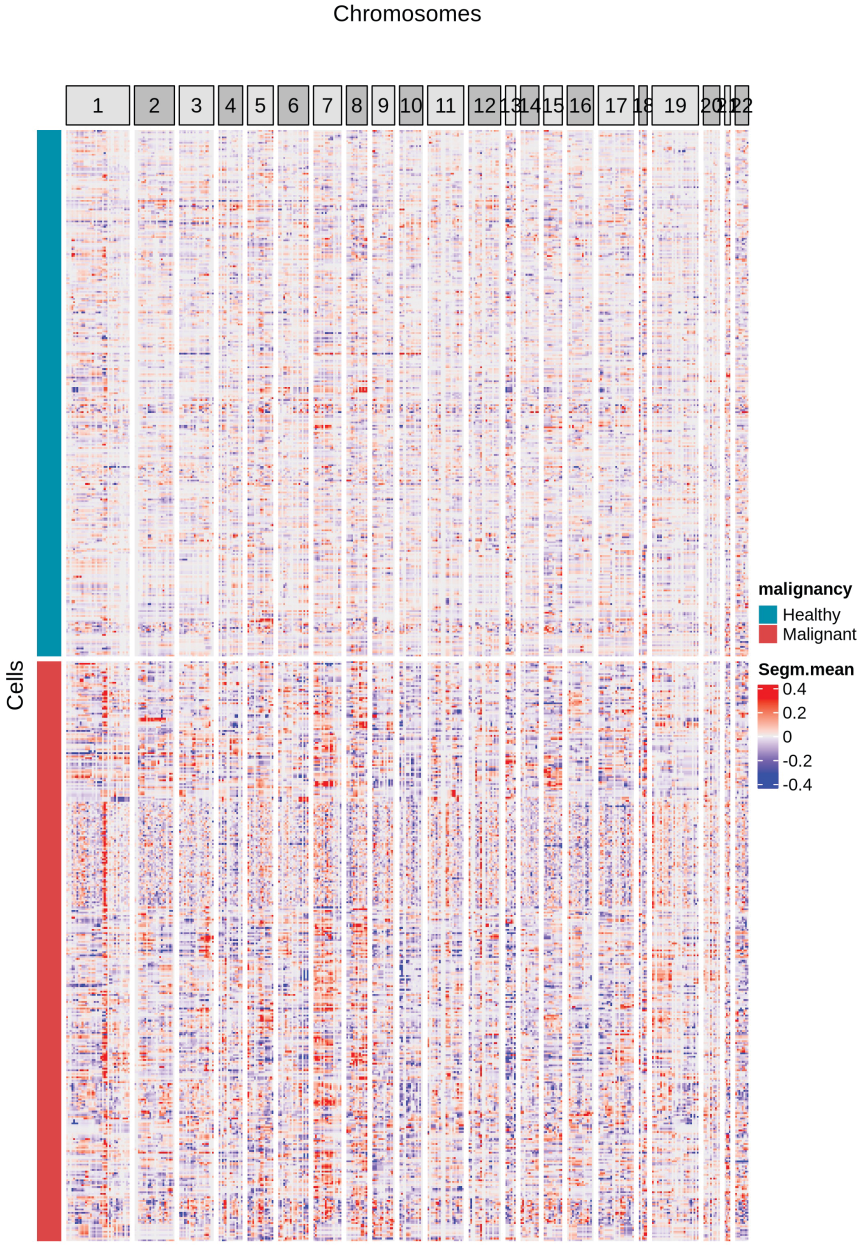


**Supplementary Figure 1. Genome-wide copy number variations (CNVs) inferred from gene expression with SCEVAN.** Each row represents a single cell, and each column represents a genomic segment grouped by chromosome. Segment mean values ($log_2$ ratio of observed copy number relative to diploid reference) are shown using a blue-to-red gradient, with blue indicating deletions (negative values), white near zero indicating neutral copy number, and red indicating amplifications (positive values). CNAs successfully inferred in 95% of samples of the cohort (810/853 samples). CNV inference from gene expression profiles separates malignant cells with CNAs (bottom) from non-malignant cells without CNAs (top).

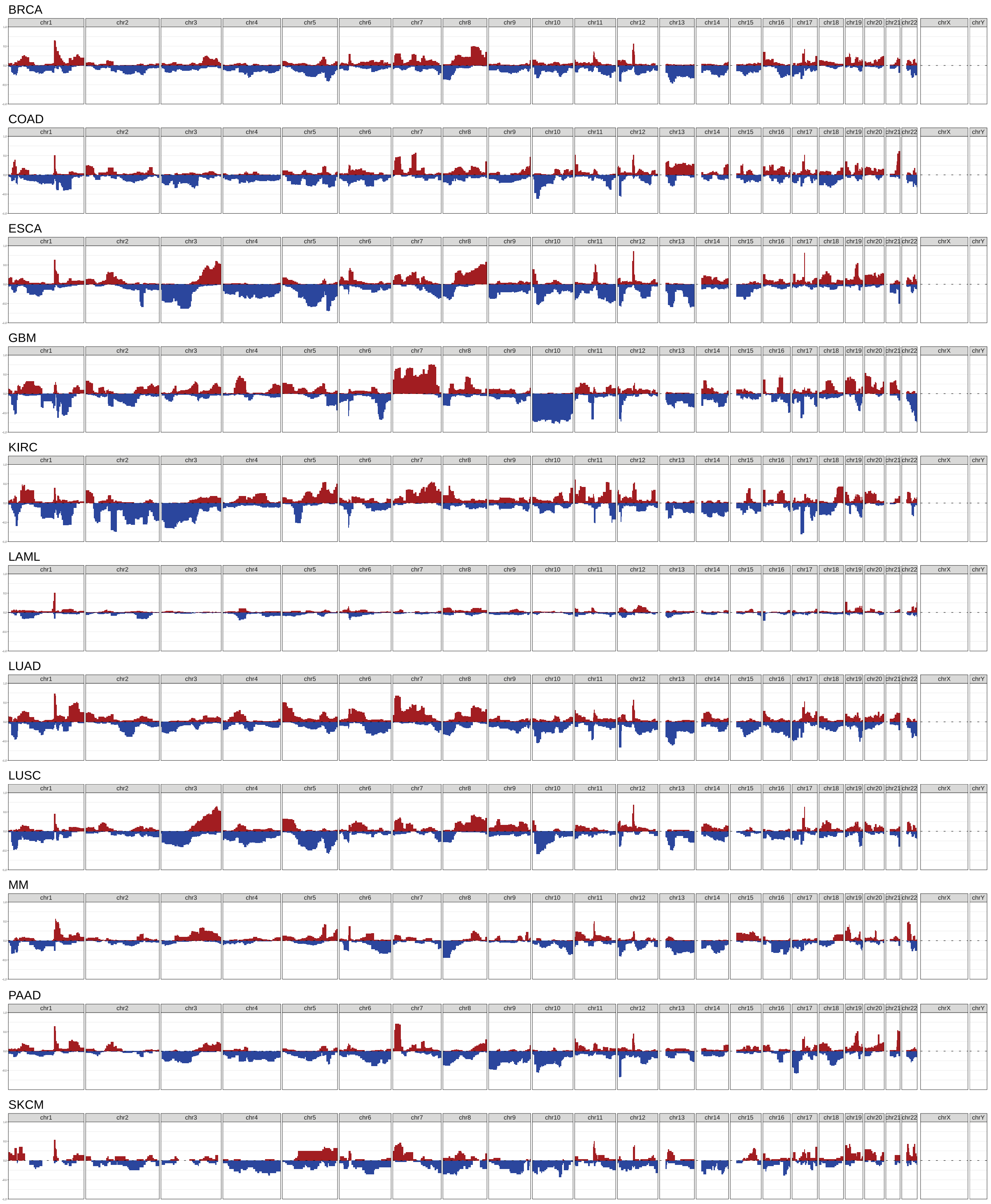


**Supplementary Figure 2. CNV landscapes in TCCA samples across the most prevalent cancer types.** The y-axis represents the proportion of subclones within each cancer type harboring CNVs, and the x-axis corresponds to chromosomal positions.

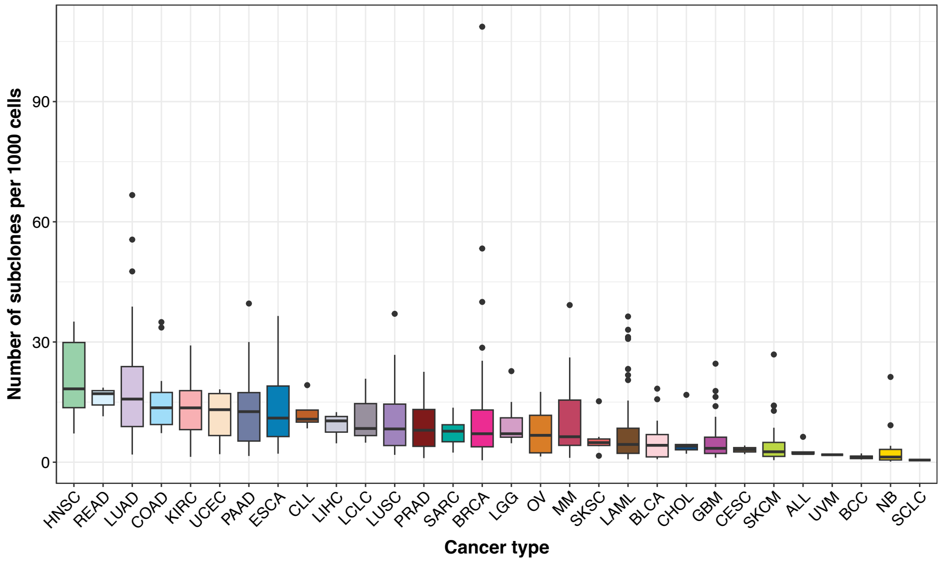


**Supplementary Figure 3. Subclonal diversity across cancer types.** Number of subclones per 1,000 cells, where subclones are defined by grouping cells with similar CNV profiles inferred by SCEVAN. Each point represents one patient sample (cell line samples excluded, so the plot includes 631 of the 810 samples with successfully inferred CNVs and subclones). Cancer types are ordered by the median number of subclones per 1,000 cells.

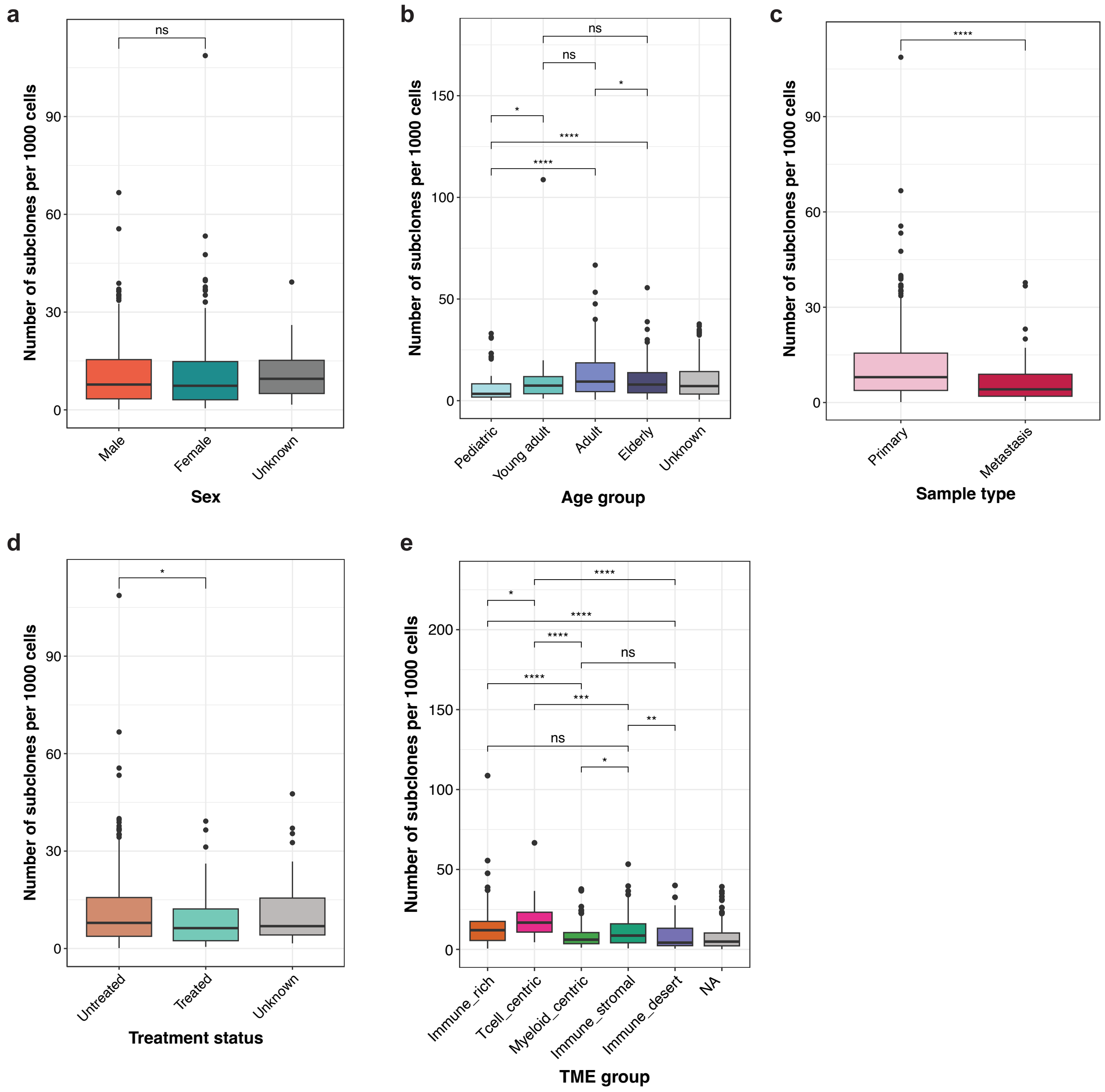


**Supplementary Figure 4. Subclonal diversity across clinical variables and TME archetypes.** Number of subclones per 1,000 cells is shown for samples grouped by (**a**) sex, (**b**) age groups (Pediatric 0–15, Young adult 16–39, Adult 40–64, Elderly ≥65), (**c**) sample type (primary vs. metastasis), (**d**) treatment status (treated vs. untreated), and (**e**) TME archetype (NA = unclassified samples, e.g., liquid tumors or samples without enough immune/stromal cells; n = 150). Group comparisons were performed using Wilcoxon tests with significance indicated as: **** p ≤ 0.0001, *** p ≤ 0.001, ** p ≤ 0.01, * p ≤ 0.05, ns p > 0.05.

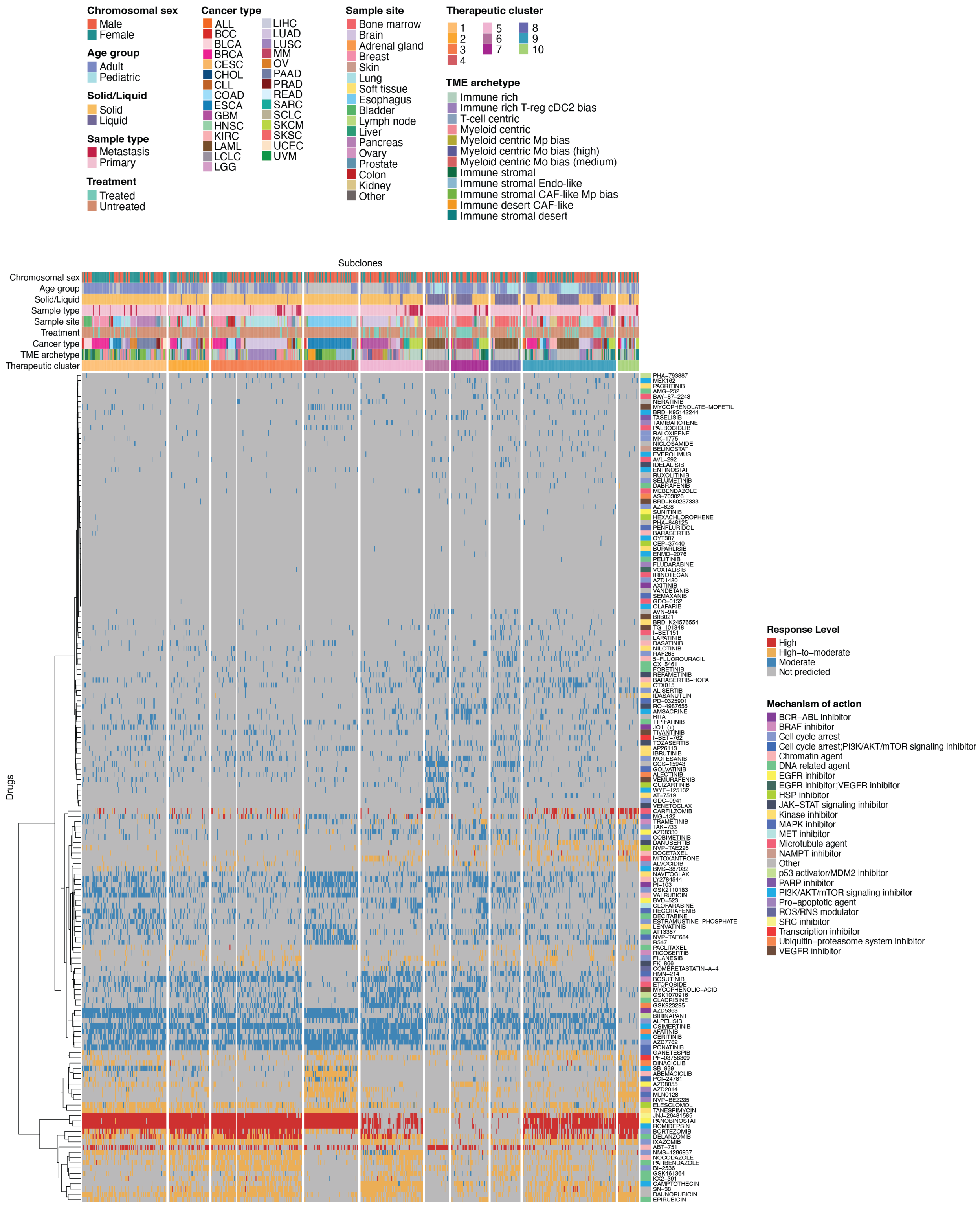


**Supplementary Figure 5. scTherapy predictions across 29 cancer types in the TCCA cohort.** Heatmap showing scTherapy-predicted therapies across patient genomic subclones from multiple tumor types. Each column corresponds to a genomic subclone (only high-confidence malignant cells were used: 611 samples, 2,627 subclones, 637,062 cells; see Methods). Annotations include sex, age, sample site, primary vs. metastasis status, solid vs. liquid tumor type, treatment condition, cancer type, TME archetype, and therapeutic cluster defined by spectral clustering of pairwise Jaccard similarities between subclone drug-prediction sets. Rows represent therapies, color-coded by mechanism of action, and the heatmap colors indicate the predicted response level. Cases where scTherapy did not return a prediction for a given genomic subclone are indicated in gray.

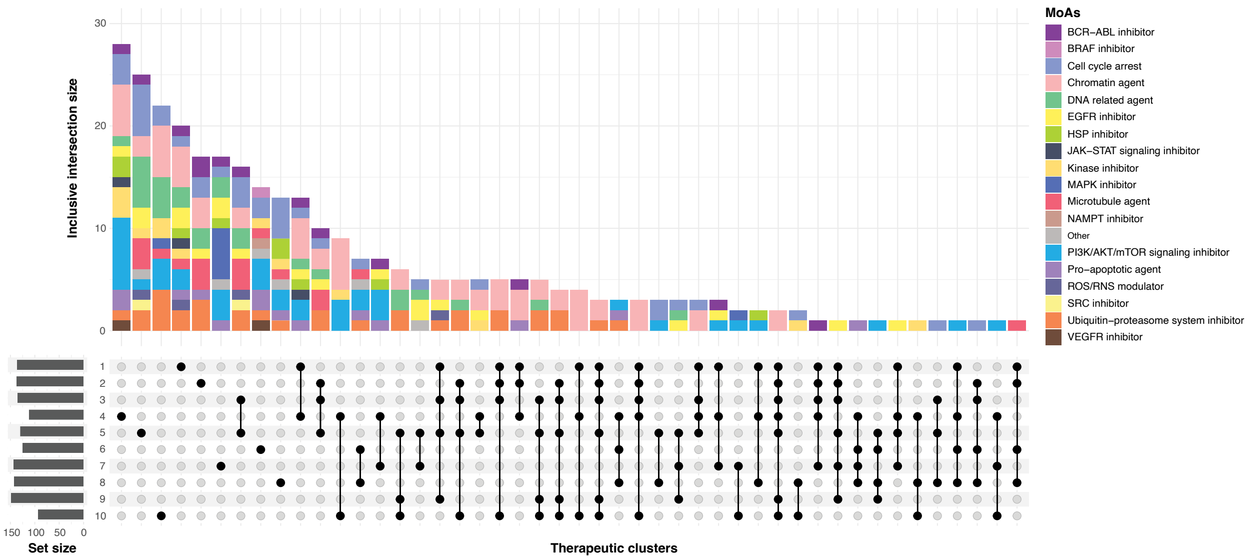


**Supplementary Figure 6. Drugs significantly associated with therapeutic clusters.** UpSet plot showing drugs enriched in one or more TCs. Each column represents a TC combination, with vertical bars showing the number of drugs in that inclusive intersection (includes all drugs present in the defining clusters, even if they also appear in other clusters). Bars are colored by the drug's mechanism of action (MoA). Only drugs predicted to be effective in ≥40% of subclones within a TC and with FDR-adjusted p-value ≤ 0.05 are shown. Associations were tested using a Chi-squared test; if any expected count was <5, a Fisher's exact test was used instead. P-values were adjusted with the Benjamini-Hochberg method. Horizontal bars on the left show the total number of drugs predicted for each TC. If a TC has no unique drugs—all predicted drugs also appear in other clusters—the inclusive intersection is not displayed (e.g., cluster 9).

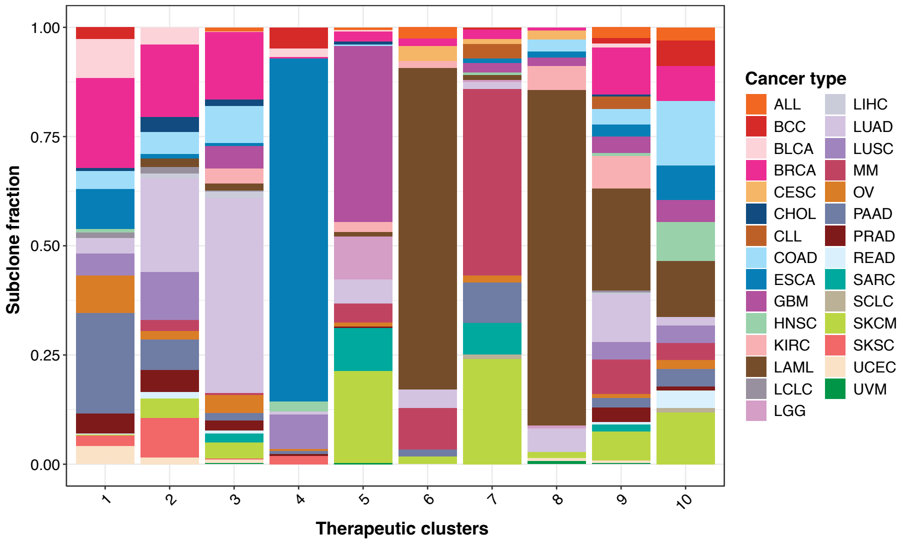


**Supplementary Figure 7. Distribution of cancer types across therapeutic clusters.** Bar plot showing the fraction of subclones from each cancer type (y-axis) within each therapeutic cluster (x-axis).

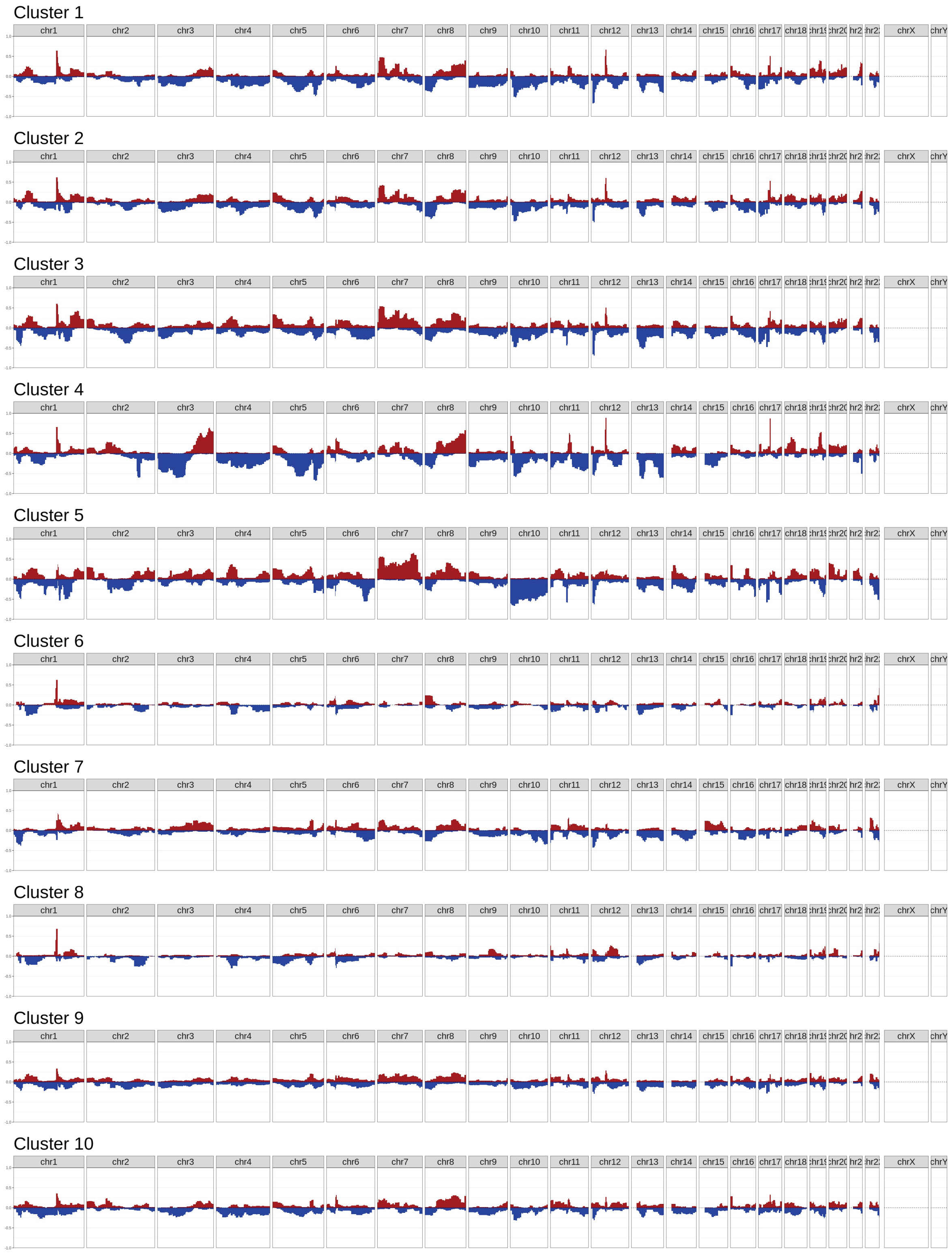


**Supplementary Figure 8. CNV landscapes across** therapeutic **clusters.** The y-axis represents the proportion of subclones within each thearpeutic cluster harboring CNVs, and the x-axis corresponds to chromosomal positions.

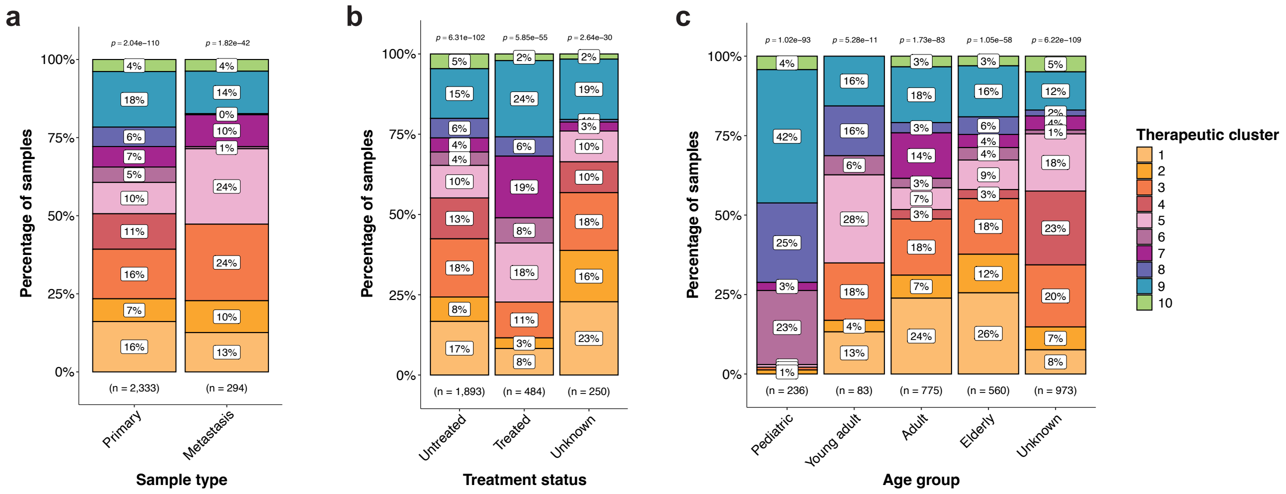


**Supplementary Figure 9. Distribution of the 10 therapeutic clusters across clinical variables. a** Sample type (primary vs metastatic), and **b** treatment status (treated vs untreated), **c** age group (Pediatric: 0–15, Young adult: 16–39, Adult: 40–64, Elderly: ≥65). The “Unknown” category indicates subclones from samples without available clinical information. Colors represent different TCs. Numbers below each bar indicate the total number of subclones per category, and numbers above show the p-value from a Chi-squared test of independence, testing whether TCs distributions are independent of the clinical variable (a low p-value suggests a biased, non-random distribution).

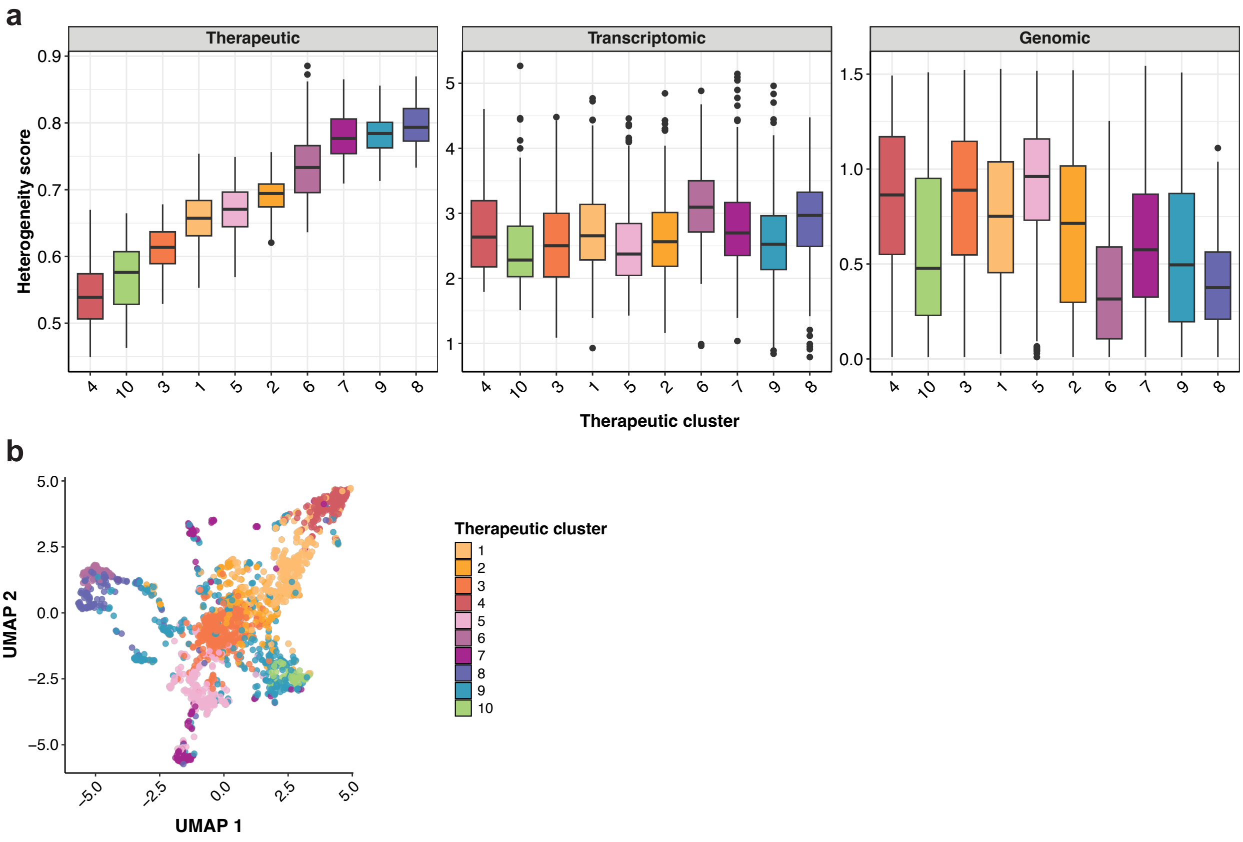


**Supplementary Figure 10. Quantitative analysis of therapeutic, transcriptomic, and genomic heterogeneity within subclones. a** Boxplots showing heterogeneity scores for each subclone across therapeutic clusters. Each point represents a subclone. Transcriptomic heterogeneity was measured as the mean covariance of gene expression (using only HVGs) across cells within each subclone, while genomic heterogeneity were measured using Shannon entropy of copy number profiles per subclone. Therapeutic heterogeneity for each subclone was calculated as the mean Jaccard distance between its predicted drug profile and all other subclones in the same cluster. Therapeutic clusters on the x-axis are ordered from lowest to highest median therapeutic heterogeneity. **b** UMAP visualization of subclones based on pairwise Jaccard distances of their drug predictions. Points are colored by therapeutic cluster assignment.

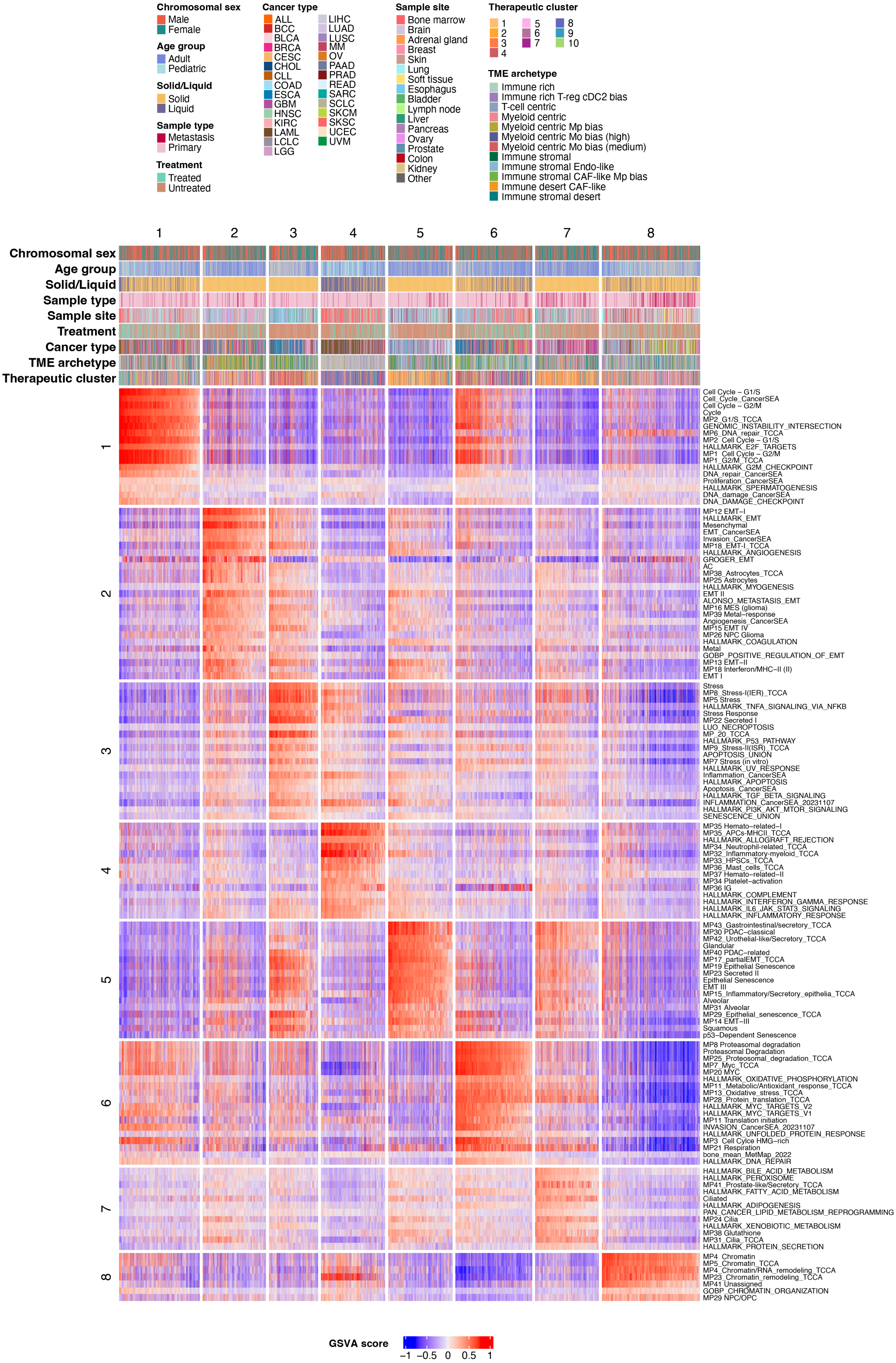
Chromosomal sex
Male
Female
Age group
Adult
Pediatric
Solid/Liquid
Solid
Liquid
Sample type
Metastasis
Primary
Treatment
Treated
Untreated
Cancer type
ALL
BCC
BLCA
BRCA
CESC
CHOL
CLL
COAD
ESCA
GBM
HNSC
KIRC
LAML
LCLC
LGG
LIHC
LUAD
LUSC
MM
OV
PAAD
PRAD
READ
SARC
SCLC
SKCM
SKSC
UCEC
UVM
Sample site
Bone marrow
Brain
Adrenal gland
Breast
Skin
Lung
Soft tissue
Esophagus
Bladder
Lymph node
Liver
Pancreas
Ovary
Prostate
Colon
Kidney
Other
Therapeutic cluster
1
2
3
4
5
6
7
8
9
10
TME archetype
Immune rich
Immune rich T-reg cDC2 bias
T-cell centric
Myeloid centric
Myeloid centric Mp bias
Myeloid centric Mo bias (high)
Myeloid centric Mo bias (medium)
Immune stromal
Immune stromal Endo-like
Immune stromal CAF-like Mp bias
Immune desert CAF-like
Immune stromal desert
1 2 3 4 5 6 7 8
Chromosomal sex
Age group
Solid/Liquid
Sample type
Sample site
Treatment
Cancer type
TME archetype
Therapeutic cluster
1
Cell Cycle – G1/S
Cell_Cycle_CancerSEA
Cell Cycle – G2/M
Cycle
MP2_G1/S_TCCA
GENOMIC_INSTABILITY_INTERSECTION
MP6_DNA_repair_TCCA
MP2 Cell Cycle – G1/S
HALLMARK_E2F_TARGETS
MP1 Cell Cycle – G2/M
MP1_G2/M_TCCA
HALLMARK_G2M_CHECKPOINT
DNA_repair_CancerSEA
Proliferation_CancerSEA
HALLMARK_SPERMATOGENESIS
DNA_damage_CancerSEA
DNA_DAMAGE_CHECKPOINT
2
MP12 EMT–I
HALLMARK_EMT
Mesenchymal
EMT_CancerSEA
Invasion_CancerSEA
MP18_EMT-I_TCCA
HALLMARK_ANGIOGENESIS
GROGER_EMT
AC
MP38_Astrocytes_TCCA
MP25 Astrocytes
HALLMARK_MYOGENESIS
EMT II
ALONSO_METASTASIS_EMT
MP16 MES (glioma)
MP39 Metal–response
Angiogenesis_CancerSEA
MP15 EMT IV
MP26 NPC Glioma
HALLMARK_COAGULATION
Metal
GOBP_POSITIVE_REGULATION_OF_EMT
MP13 EMT–II
MP18 Interferon/MHC–II (II)
EMT I
3
Stress
MP8_Stress-I(IER)_TCCA
MP5 Stress
HALLMARK_TNFA_SIGNALING_VIA_NFKB
Stress Response
MP22 Secreted I
LUO_NECROPTOSIS
MP_20_TCCA
HALLMARK_P53_PATHWAY
MP9_Stress-II(ISR)_TCCA
APOPTOSIS_UNION
MP7 Stress (in vitro)
HALLMARK_UV_RESPONSE
Inflammation_CancerSEA
HALLMARK_APOPTOSIS
Apoptosis_CancerSEA
HALLMARK_TGF_BETA_SIGNALING
INFLAMMATION_CancerSEA_20231107
HALLMARK_PI3K_AKT_MTOR_SIGNALING
SENESCENCE_UNION
4
MP35 Hemato–related–I
MP35_APCs-MHCII_TCCA
HALLMARK_ALLOGRAFT_REJECTION
MP34_Neutrophil-related_TCCA
MP32_Inflammatory-myeloid_TCCA
MP33_HPSCs_TCCA
MP36_Mast_cells_TCCA
MP37 Hemato–related–II
MP34 Platelet–activation
MP36 IG
HALLMARK_COMPLEMENT
HALLMARK_INTERFERON_GAMMA_RESPONSE
HALLMARK_IL6_JAK_STAT3_SIGNALING
HALLMARK_INFLAMMATORY_RESPONSE
5
MP43_Gastrointestinal/secretory_TCCA
MP30 PDAC–classical
MP42_Urothelial-like/Secretory_TCCA
Glandular
MP40 PDAC–related
MP17_partialEMT_TCCA
MP19 Epithelial Senescence
MP23 Secreted II
Epithelial Senescence
EMT III
MP15_Inflammatory/Secretory_epithelia_TCCA
Alveolar
MP31 Alveolar
MP29_Epithelial_senescence_TCCA
MP14 EMT–III
Squamous
p53–Dependent Senescence
6
MP8 Proteasomal degradation
Proteasomal Degradation
MP25_Proteosomal_degradation_TCCA
MP7_Myc_TCCA
MP20 MYC
HALLMARK_OXIDATIVE_PHOSPHORYLATION
MP11_Metabolic/Antioxidant_response_TCCA
MP13_Oxidative_stress_TCCA
MP28_Protein_translation_TCCA
HALLMARK_MYC_TARGETS_V2
HALLMARK_MYC_TARGETS_V1
MP11 Translation initiation
INVASION_CancerSEA_20231107
HALLMARK_UNFOLDED_PROTEIN_RESPONSE
MP3 Cell Cylce HMG–rich
MP21 Respiration
bone_mean_MetMap_2022
HALLMARK_DNA_REPAIR
7
HALLMARK_BILE_ACID_METABOLISM
HALLMARK_PEROXISOME
MP41_Prostate-like/Secretory_TCCA
HALLMARK_FATTY_ACID_METABOLISM
Ciliated
HALLMARK_ADIPOGENESIS
PAN_CANCER_LIPID_METABOLISM_REPROGRAMMING
MP24 Cilia
HALLMARK_XENOBIOTIC_METABOLISM
MP38 Glutathione
MP31_Cilia_TCCA
HALLMARK_PROTEIN_SECRETION
8
MP4 Chromatin
MP5_Chromatin_TCCA
MP4_Chromatin/RNA_remodeling_TCCA
MP23_Chromatin_remodeling_TCCA
MP41 Unassigned
GOBP_CHROMATIN_ORGANIZATION
MP29 NPC/OPC
GSVA score
−1 −0.5 0 0.5 1

**Supplementary Figure 11. Functional characterization of therapeutic clusters using own-derived MPs and external functional gene sets.** Heatmap showing GVSA scores for a representative subset of 2,533 subclone pseudobulks (columns), grouped by therapeutic cluster. Rows represent enrichment scores for 130 functional gene sets derived from internal MPs and external datasets (MSigDB Hallmarks; Barkley et al., 2022; Gavish et al., 2023). Only samples and gene sets that could be biclustered using the FABIA algorithm are shown. Top annotations indicate sample metadata: sex, age, cancer type (solid/liquid), sample type (primary/metastasis), sample site, tumor type, treatment status (treated/untreated), TME archetype, and therapeutic cluster assignment.

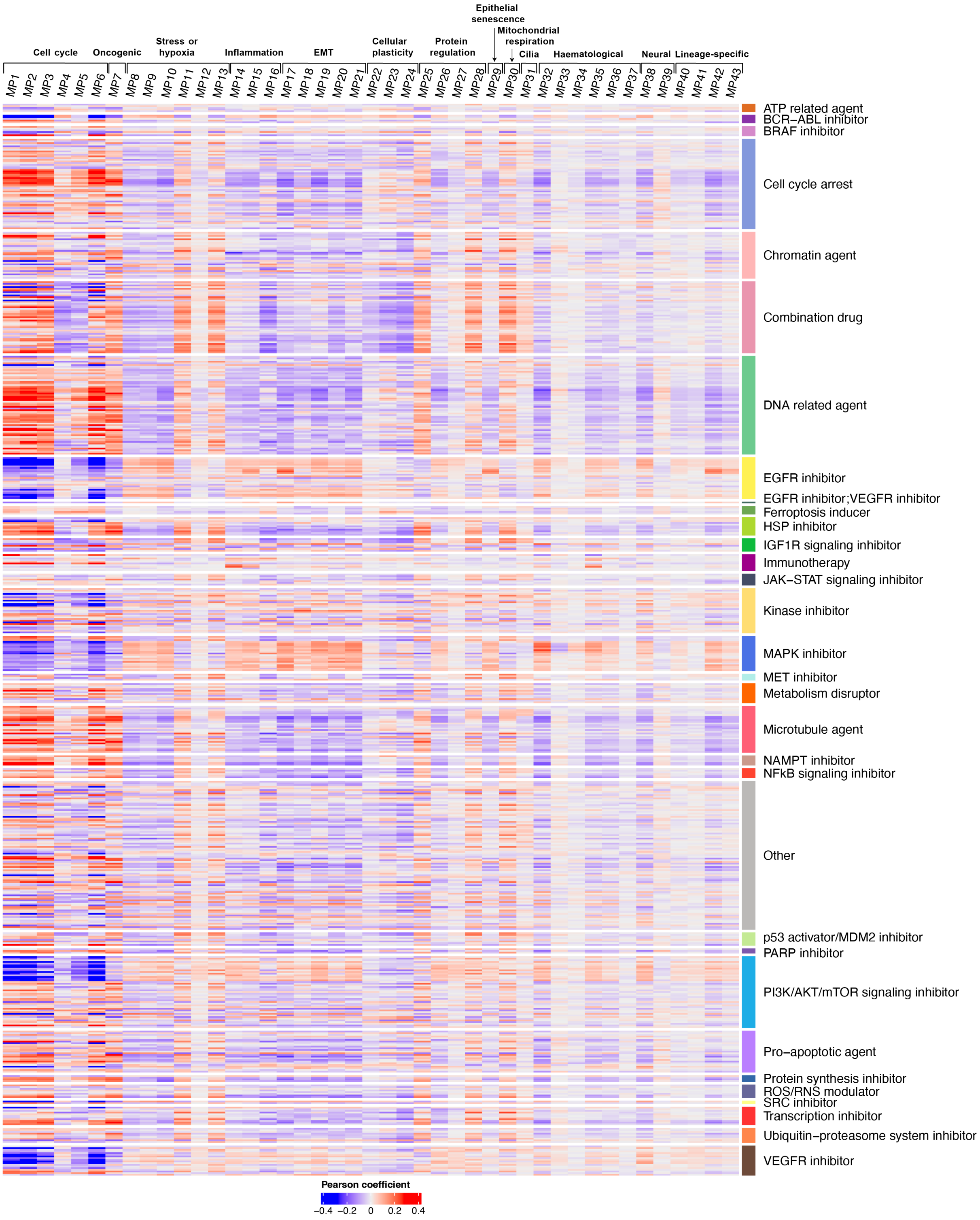

**Supplementary Figure 12. Correlation between functional metaprograms and drug sensitivities from Beyondcell.** Heatmap showing Pearson correlations between 43 functional MPs (columns) and 589 drugs from the Beyondcell drug sensitivity collection, including immunotherapies (SSc; rows). Drug names are omitted due to the large number of compounds; rows are instead color-coded and labeled by their mechanism of action.

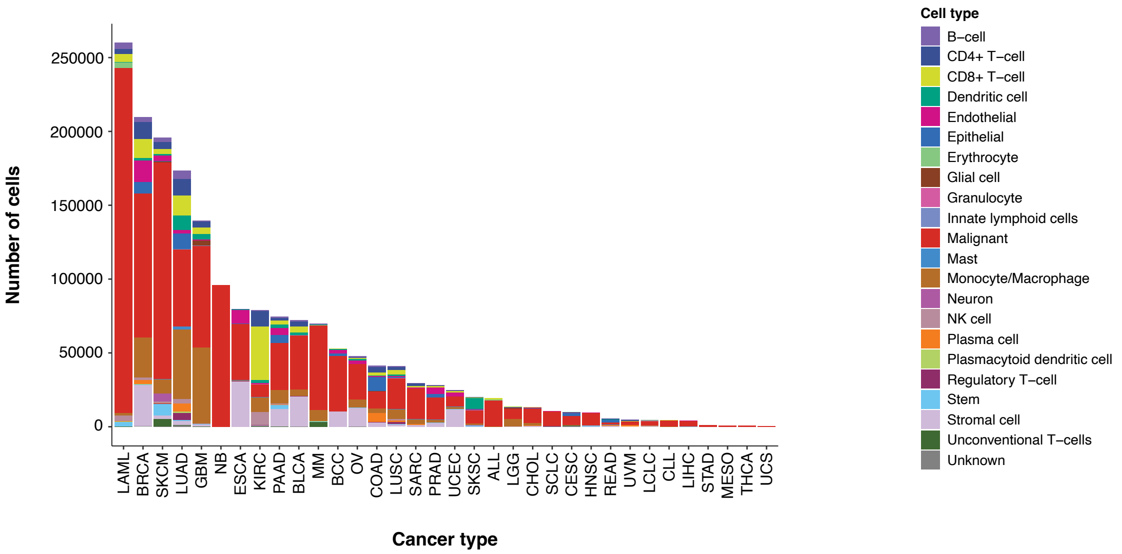


**Supplementary Figure 13. Distribution of cell types across cancer types.** Each stacked bar shows the number of cells for each cell type within each cancer type, with cancer types arranged from highest to lowest total cell count. Colors indicate the different cell types, enabling comparison of cell type composition across cancers.

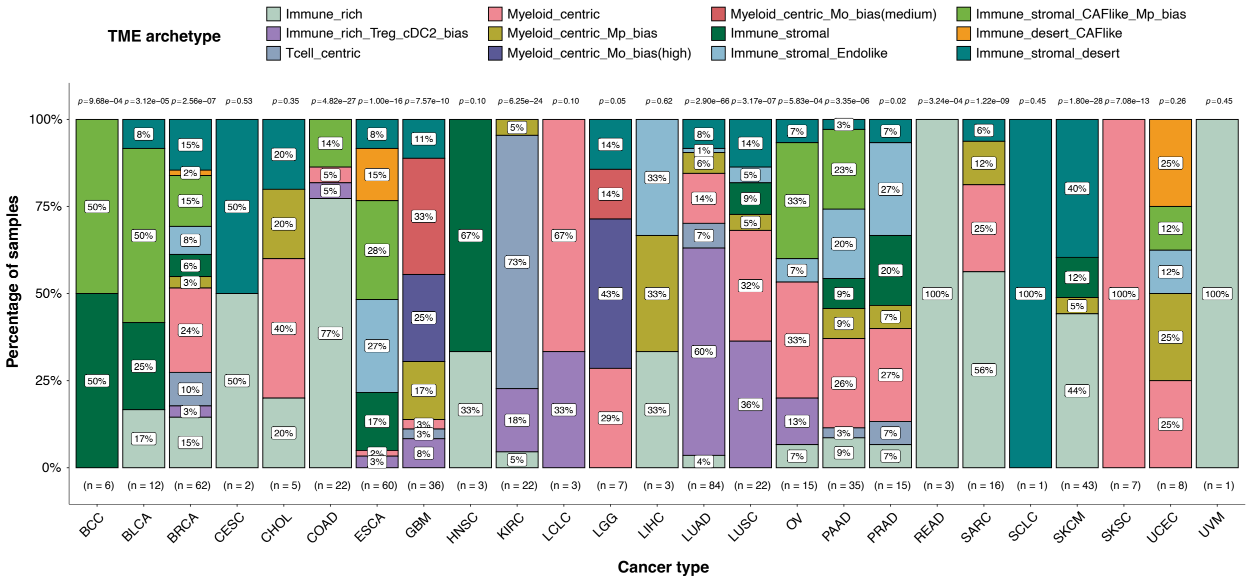


**Supplementary Figure 14. Distribution of the 12 TME archetypes across cancer types.** Stacked bars show the percentage of samples assigned to each TME archetype within each cancer type. Only 493 of 853 samples could be classified (the remaining samples are liquid tumors or lack sufficient immune and stromal cells). Colors indicate different TME archetypes, facilitating comparison of TME composition across cancer types. Numbers below each bar indicate the total number of samples classified in that cancer type, and numbers above each bar show the p-value from a Chi-squared test of independence, indicating whether the overall distribution of TME archetypes differs across cancer types (a low p-value suggests a biased, non-random distribution).

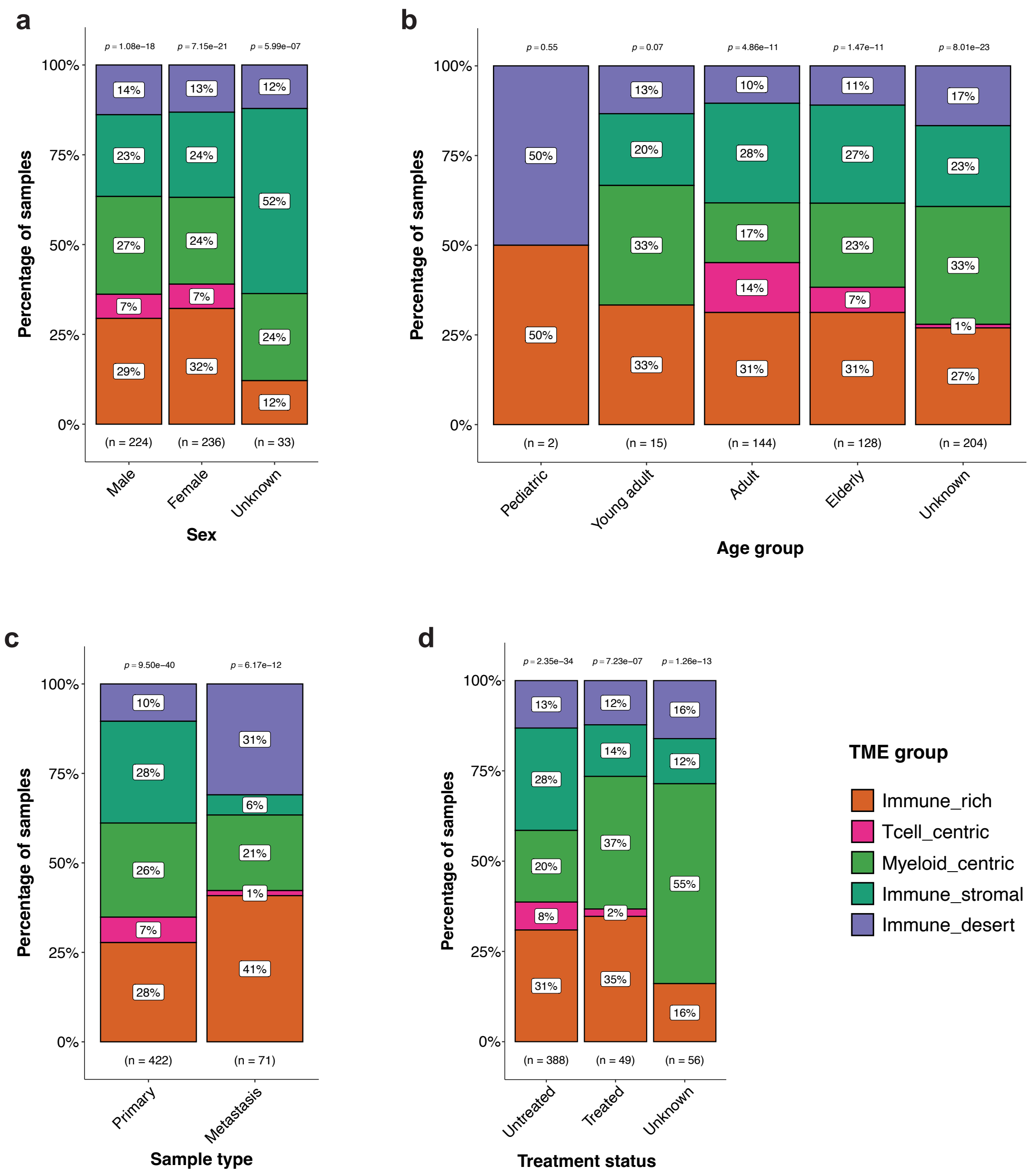


**Supplementary Figure 15. Distribution of the five general TME groups across clinical variables.** **a** Sex, **b** age group (Pediatric: 0–15, Young adult: 16–39, Adult: 40–64, Elderly: ≥65), **c** sample type (primary vs metastatic), and **d** treatment status (treated vs untreated). The “Unknown” category indicates samples without available clinical information. Colors represent different TME groups. Numbers below each bar indicate the total number of classified samples per category, and numbers above show the p-value from a Chi-squared test of independence, testing whether TME distributions are independent of the clinical variable (a low p-value suggests a biased, non-random distribution).

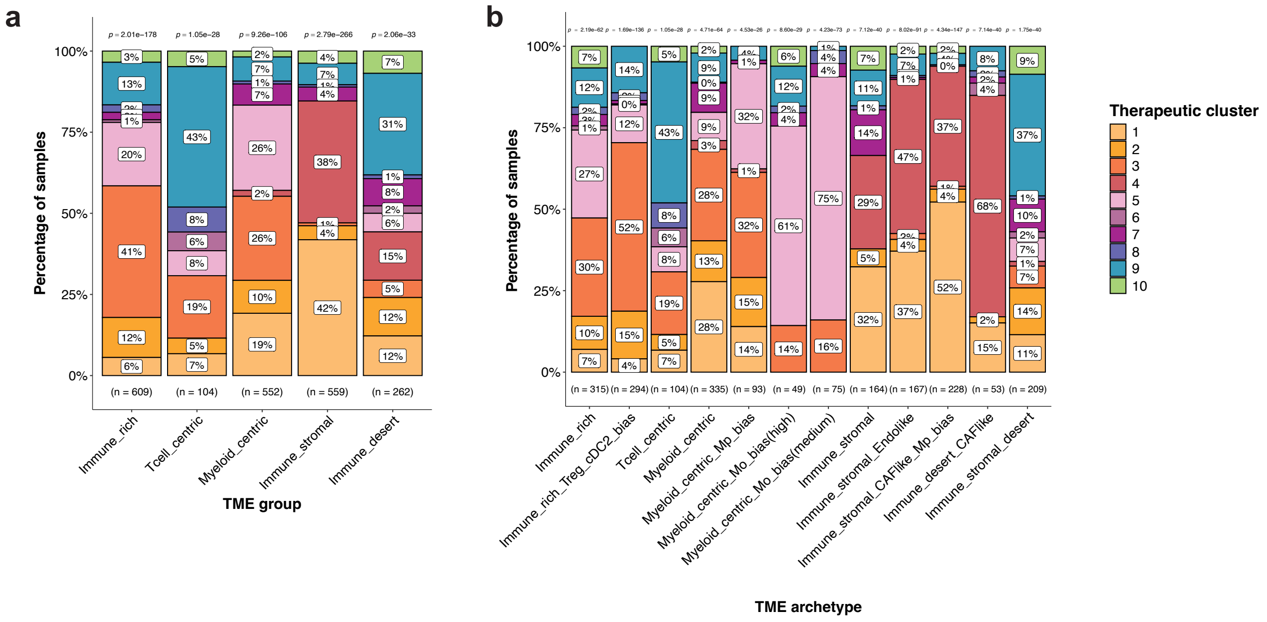


**Supplementary Figure 16. Distribution of the 10 therapeutic clusters (TCs) across TME archetypes. a** TME archetype groups. **b** Individual TME archetypes. Only subclones from samples with TME annotations are included (n = 2,086 subclones from 480 samples; the "none" category was excluded). Bars are colored by TC. Numbers below each bar indicate the total number of subclones in each category, and numbers above bars show the p-value from a Chi-squared test of independence, assessing whether TC distributions are independent of the TME archetype(low p-values indicate non-random, biased distributions).

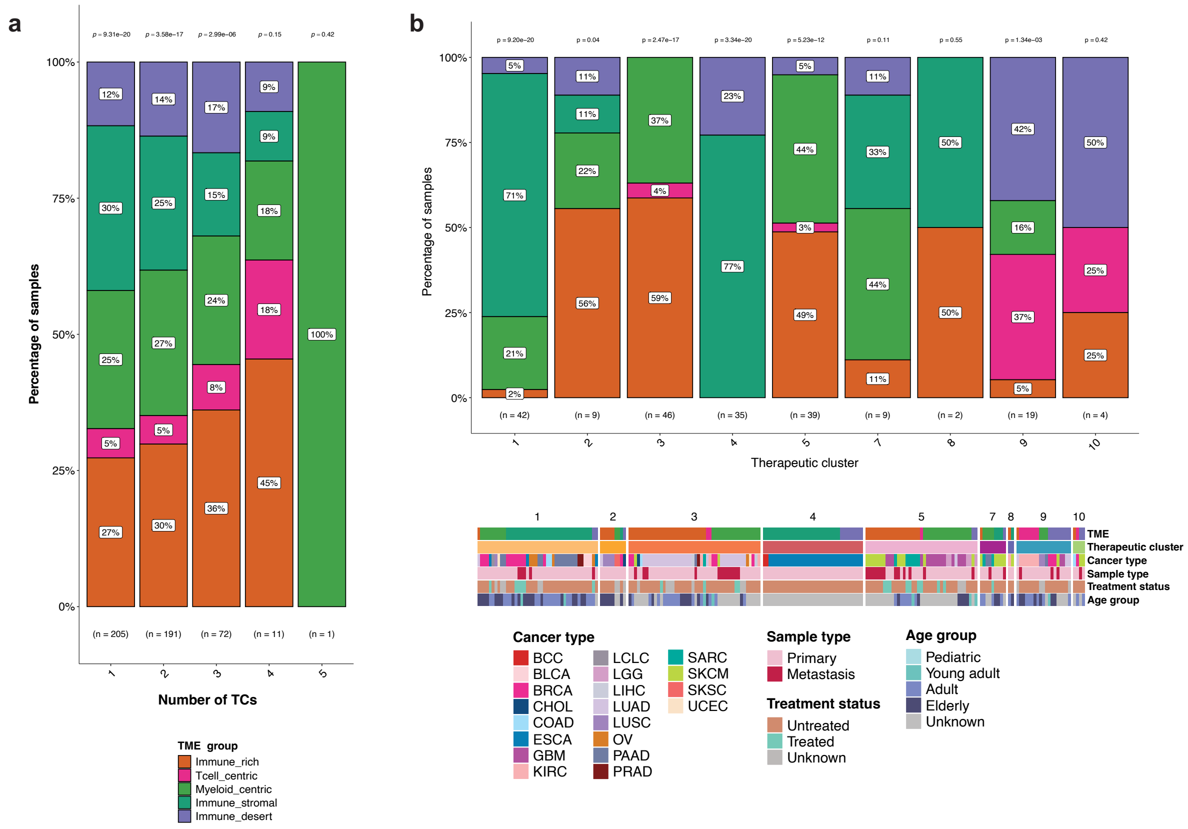


**Supplementary Figure 17. Distribution of TME archetype groups across therapeutic clusters (TCs), considering only samples with TME annotation. A** Proportion of samples assigned to 1, 2, 3, 4, or 5 TCs, colored by their TME archetype group. **B** For samples where all subclones fall into a single TC (first bar in panel A, n = 205), one-to-one relationships between sample, TC, TME archetype, and clinical information are shown. Bars represent the proportion of samples of each TME archetype group within each TC. Below, additional sample-level clinical variables are displayed, including cancer type, sample type (primary/metastasis), treatment status (treated/untreated), and age group (Pediatric: 0–15, Young adult: 16–39, Adult: 40–64, Elderly: ≥65).

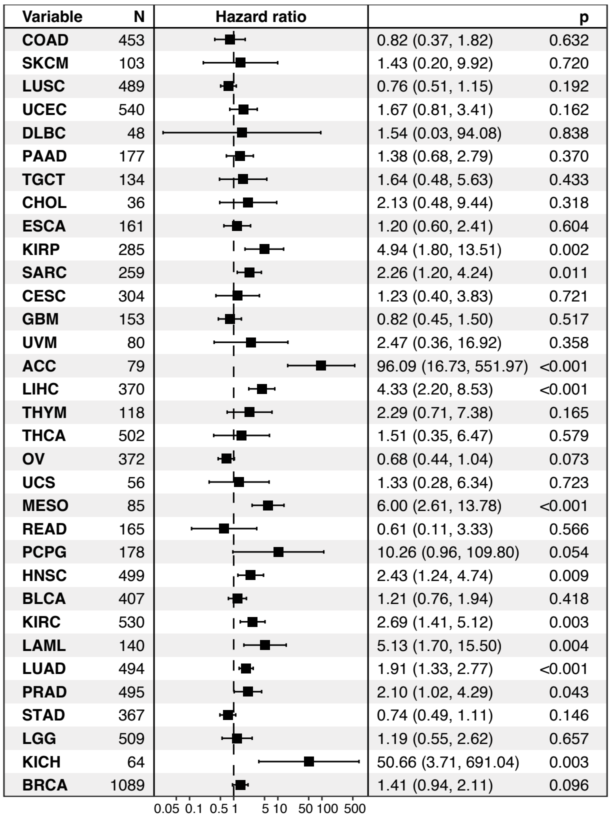


**Supplementary Figure 18. Expression of therapeutic cluster 10 markers in TCGA and their association with patient survival.** Forest plot showing hazard ratios and p-values from Cox proportional hazards models (adjusted for age and sex) assessing the relationship between TC10 signature and survival across 33 TCGA cancer types.

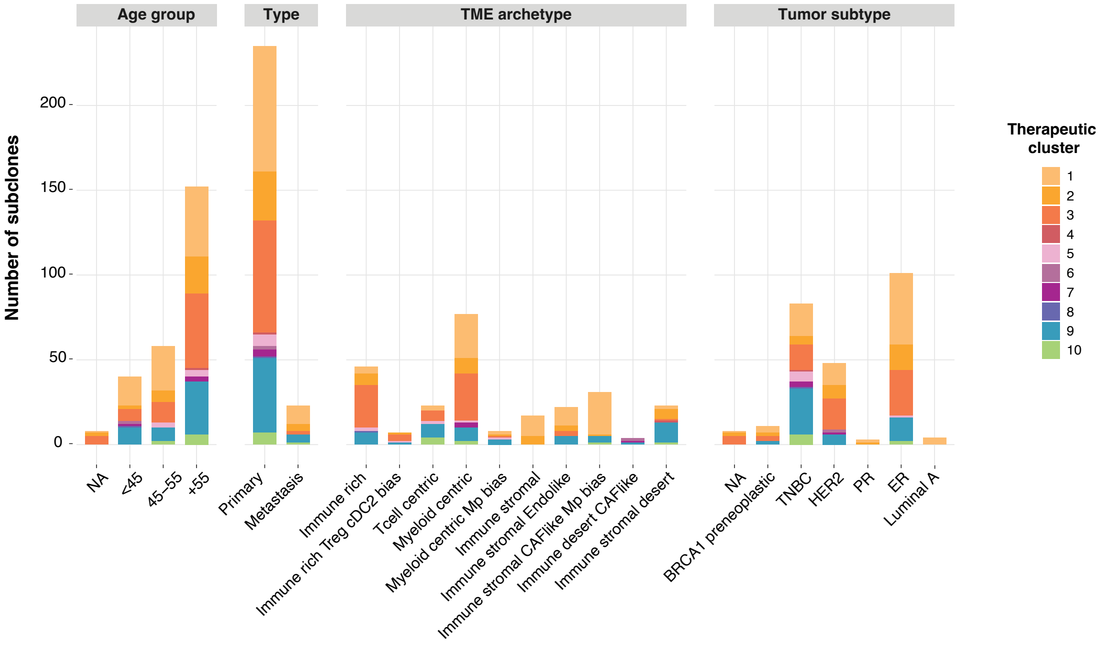


**Supplementary Figure 19. Distribution of therapeutic clusters across clinical variables in breast cancer subclones.** Clinical variables include age group, primary versus metastatic status, tumor microenvironment (TME) archetype, and molecular subtype. Stacked bar plots show the proportion of subclones assigned to each therapeutic cluster. Age groups are defined as <45, 45–55, and >55 years and are used as an age-based proxy for menopausal status. Molecular subtypes include preneoplastic, triple-negative breast cancer (TNBC), HER2-positive, progesterone receptor–positive (PR), estrogen receptor–positive (ER), and Luminal A.

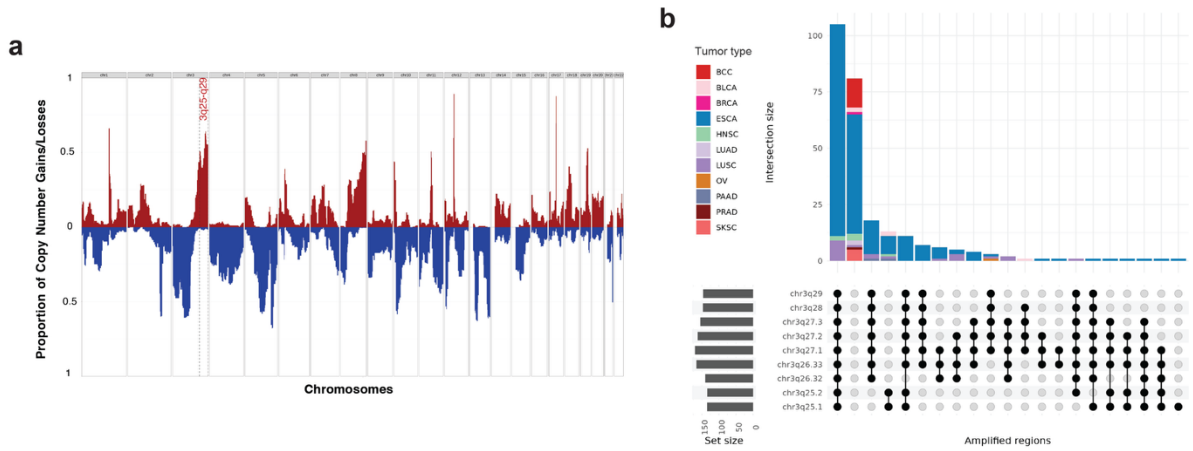


**Supplementary Figure 20. Distinct 3q25–q29 copy-number amplification and subclonal CNV combinations in TC4. a** CNV profile of therapeutic cluster 4 (TC4), highlighting the 3q25–q29 genomic region, which is uniquely amplified in this cluster compared to all others. **b** UpSet plot showing combinations of copy-number amplifications in TC4 subclones. Vertical bars represent the number of subclones carrying each combination, colored by cancer type. Horizontal bars indicate the total number of TC4 subclones with each individual amplification.

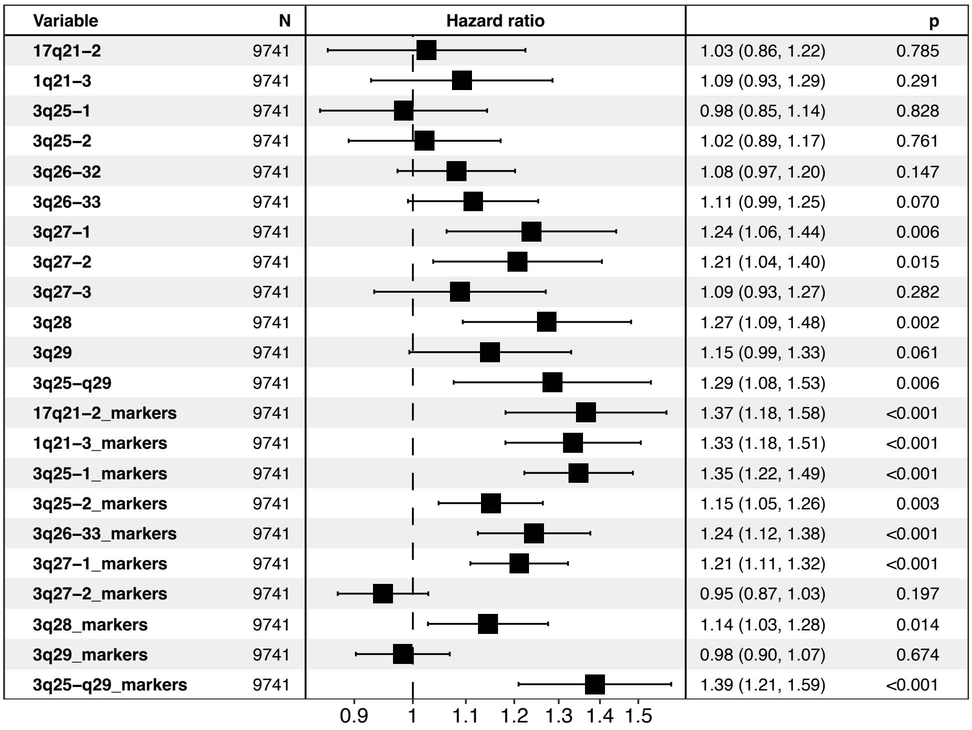


**Supplementary Figure 21. Prognostic associations of genomic band–derived and TC4-marker gene signatures.** Forest plot showing hazard ratios and p-values from Cox models (adjusted for age, sex, clinical stage, and tumor grade) evaluating gene signatures derived from (i) individual genomic bands and (ii) the intersection between these bands and TC4 marker genes.

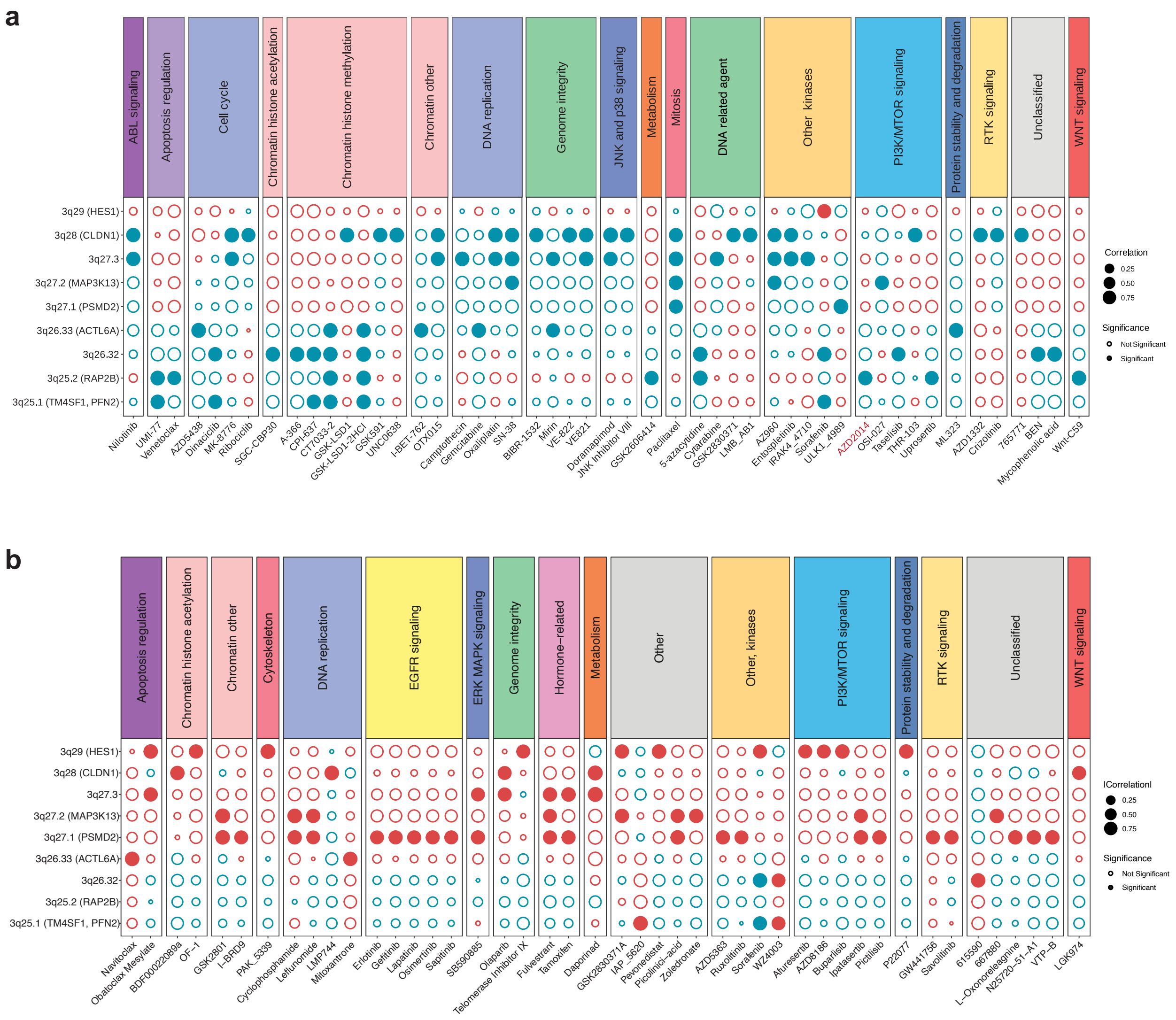


**Supplementary Figure 22. Correlation between TC4-enriched genomic band signatures (>50% frequency in TC4 subclones) and drug response in ESCA cell lines from the GDSC dataset.** On the y-axis, band-derived gene signatures are shown, with representative marker genes indicated in parentheses when overlapping with TC4 markers (see Fig. 2c). On the x-axis, GDSC drugs are grouped and color-coded by mechanism of action. **a** Only drugs with at least one negative correlation are displayed; negative correlations indicate that higher genomic band signature expression is associated with lower AUC values, reflecting greater drug sensitivity. **b** Only drugs with at least one positive correlation are displayed; positive correlations indicate that higher genomic band signature expression is associated with higher AUC values, reflecting lower drug sensitivity.
